# Sweet revenge - *Streptococcus pyogenes* showcases first example of immune evasion through specific IgG glycan hydrolysis

**DOI:** 10.1101/544064

**Authors:** Andreas Naegeli, Eleni Bratanis, Christofer Karlsson, Oonagh Shannon, Raja Kalluru, Adam Linder, Johan Malmström, Mattias Collin

## Abstract

*Streptococcus pyogenes* (Group A streptococcus, GAS) is an important human pathogen responsible for a wide variety of diseases from uncomplicated tonsillitis to life-threatening invasive infections. GAS secretes EndoS, an endoglycosidase able to specifically cleave the conserved *N*-glycan on human IgG antibodies. *In vitro*, removal of this glycan impairs IgG effector functions but its relevance to GAS infection *in vivo* is unclear. Using targeted mass spectrometry, we were able to characterize the effects of EndoS on host IgG glycosylation during the course of natural infections in human patients. We found substantial IgG glycan hydrolysis locally at site of infection as well as systemically in the most severe cases. Using these findings we were able to set up appropriate model systems to demonstrate decreased resistance to phagocytic killing of GAS lacking EndoS *in vitro*, as well as decreased virulence in a mouse model of invasive infection. This study represents the first described example of specific bacterial IgG glycan hydrolysis during infection and highlights the importance of IgG glycan hydrolysis for streptococcal pathogenesis. We thereby offer new insights into the mechanism of immune evasion employed by this pathogen with clear implications for treatment of severe GAS infections and future efforts at vaccine development.

## Introduction

*Streptococcus pyogenes* (group A streptococcus, GAS) is a human pathogen causing a diverse range of diseases. GAS can cause mild infections such as tonsillitis and impetigo but also severe diseases such as streptococcal toxic shock syndrome, necrotizing fasciitis, and erysipelas^1^. Furthermore, repeated and/or untreated GAS infections can trigger serious postinfectious immune-mediated disorders, including acute poststreptococcal glomerulonephritis, acute rheumatic fever, and rheumatic heart disease^1^. With a prevalence of at least 18 million severe cases, leading to approximately half a million deaths world wide annually as well as over 700 million annual cases of mild infections^2^, GAS infections are a large public health burden.

Protective immunity towards GAS is generally poor and recurrent infections are not uncommon, especially in children^3^. This is despite the fact that most people do in fact raise an adaptive immune response and exhibit high titers of IgG antibodies towards different GAS antigens^4–7^. The reason for the lack of protection is not entirely understood but can in part be attributed to the large number of different GAS serotypes and the surface antigen variability this entails^8^. GAS is also able to counteract adaptive immunity by specifically impairing IgG function. This can be mediated by non-immune IgG binding to Fc binding proteins on the streptococcal surface such as the M and M-related proteins^9,10^ or through specific degradation of the IgGs themselves. GAS secretes for example the IgG specific protease IdeS which is able to cleave the antibody in the hinge region, separating the antigen-binding Fabs from the effector function-promoting Fc region^11^.

GAS is further able to degrade IgGs by secretion of the endoglycosidase EndoS. This enzyme cleaves the conserved Fc *N*-glycan from IgGs with great specificity^12^ (Fig. 1a). This glycan is situated at the interaction surface between the IgG and Fc receptors^13,14^ as well as the complement system^15^ and is therefore ideally located to influence IgG effector function. While an antibody’s specificity is determined by the Fab regions, the Fc region determines which effector functions are elicited and the structure of the Fc glycan has been shown to be crucial in the regulation of this process^16^. For example, IgGs lacking core fucosylation exhibit increased affinity for FcγRIIIA and are therefore significantly more potent in eliciting antibody-dependent cellular cytotoxicity^17,18^. Furthermore, the degree of galactosylation of the Fc glycan influences an IgGs ability to activate the complement system^19^. Consequently, IgG antibodies lacking the Fc glycan fail to bind to most Fc receptors and are unable to activate the complement system^20,21^. Despite the accumulating evidence that antibody glycans are instrumental in regulating and fine tuning the immune response, very little is know about the role of antibody glycosylation during infections and if it contributes to the outcome of disease. However, recent high profile studies that have suggested that the IgG glycosylation status influences whether HIV infection is controlled^22,24^ and whether *Mycobacterium tuberculosis* infection is active or latent^23^.

**FIGURE 1:**
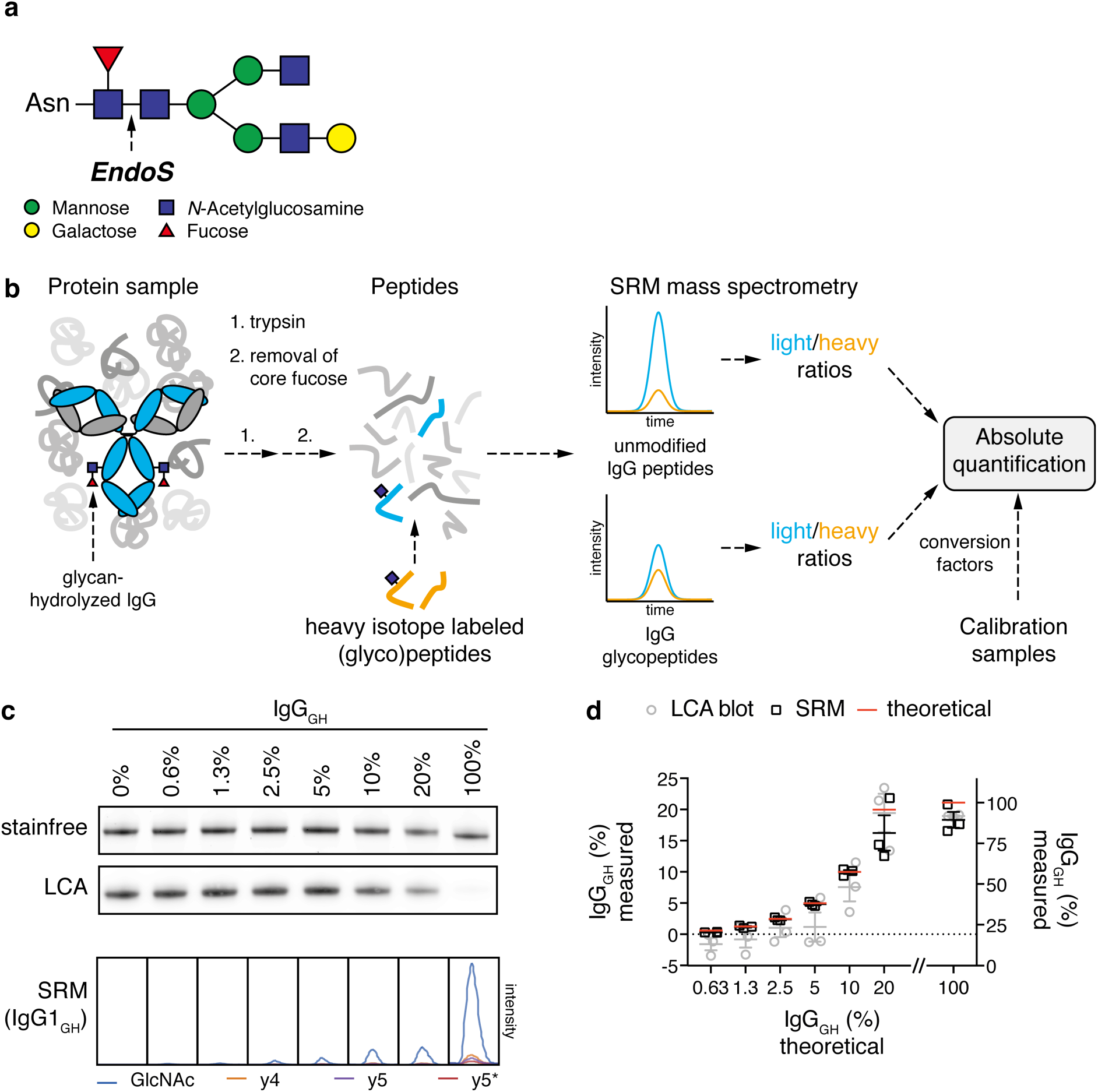
TARGETED MASS SPECTROMETRY TO QUANTIFY IGG GLYCAN HYDROLYSIS. (a) Typical *N*-glycan structure found on N297 of human IgG antibodies. The arrow marks the EndoS cleavage site in the chitobiose core of the glycan. The EndoS reaction product is an IgG carrying either a single GlcNAc or a GlcNAc-Fucose disaccharide, depending on the core fucosylation status of the antibody. (b) Overview of the SRM-MS method. Complex protein samples are digested to peptides using trypsin and potential core fucosylation is removed using α-fucosidase. The resulting peptide samples are spiked with heavy isotope labeled (glyco)peptide standards corresponding to both subclass specific IgG glycopeptides modified with a single GlcNAc residue as well as subclass specific unmodified peptides. This peptide mixture is analyzed by SRM mass spectrometry resulting in light/heavy ratios for each of the peptides of interest. The absolute amount (concentration) of each IgG subclass as well as the amount of IgG_GH_ is derived from the obtained ratios, using conversion factors determined from a defined set of standard samples c&d) Validation of SRM-MS quantitative accuracy. A set of plasma samples with defined percentages of IgG_GH_ was prepared by dilution of EndoS-treated plasma with untreated control plasma. The samples were analyzed separately in triplicate by SDS-PAGE/LCA blotting and SRM mass spectrometry. Raw data from both methods is shown in panel c. For the SRM method the chromatograms originating from the glycan-hydrolyzed IgG1 glycopeptide are shown, each transition in a different color. The asterisk denotes a fragment ion that has undergone a neutral loss of the GlcNAc modification. The degree of IgG glycan hydrolysis in the standard sample set was quantified using both methods (panel d). The red line marks the theoretical value and the measured values are depicted in grey circles (LCA blot) and black boxes (SRM) respectively. The means and standard errors are plotted and each individual data point is marked with a circle or box. The first 6 values are plotted on the left z-axis, the final 100% IgG_GH_ sample on the right y-axis.

Due to this functional importance of the Fc *N*-glycan, its hydrolysis by EndoS leads to impaired IgG effector functions such as Fc receptor binding and complement activation *in vitro*^20,24,25^. This would intuitively suggest a role for EndoS in evasion of adaptive immunity through perturbation of protective IgG responses. However, studies on the influence of EndoS on GAS pathogenesis have so far been few and inconclusive. They were unable to demonstrate under which conditions EndoS is expressed and active and had to rely on overexpression or addition of recombinant enzyme to manifest a virulence phenotype^25,26^. These efforts at elucidating the contribution of EndoS to GAS virulence have been hampered by the difficulty of finding relevant model systems and the lack of a sensitive analytical approach to quantify EndoS activity in complex systems^25,26^.

We therefore wanted to take a different approach by first characterizing the effect of EndoS on the hosts IgG glycosylation *in vivo* during the course of natural GAS infections in human patients and then use these results to set up relevant model system able to show how EndoS affects the hosts IgG response and how this contributes to GAS virulence. This necessitates an assay that is robust, specific, and sensitive enough to be able to quantify the glycosylation state of IgG in complex patient samples. We chose to employ selected reaction monitoring (SRM), a targeted mass spectrometry approach that is uniquely suitable for this type of analysis as it allows for the quantification of predefined target molecules (such as the EndoS reaction products) directly from highly complex samples. SRM is based on precursor peptide ion selection, fragmentation through collision, and detection of selected peptide fragment ions in a triple quadrupole mass spectrometer. Precursor/fragment pairs, so called transitions, are chosen that are unique to the molecule to be detected (i.e. the EndoS reaction products) and data is only acquired for these defined targets (for a review see Picotti et al.^27^). In previous studies, proteins with concentrations as low as 300 ng/ml have been reliably quantified out of crude human plasma preparations^28,29^ and SRM has also been used successfully for quantification of the different glycan structures on IgG directly from serum samples^30^.

We employed SRM-MS to quantitatively assess the EndoS-mediated IgG glycan hydrolysis in samples from natural GAS infections in humans. We used these findings to set up relevant *in vitro* assays and animal models in order to demonstrate the importance of EndoS-mediated antibody modification in evasion of adaptive immunity and therefore GAS virulence.

## Results

### A TARGETED MASS SPECTROMETRY APPROACH FOR QUANTITATIVE ANALYSIS OF IGG GLYCAN HYDROLYSIS

In order to assess the effect of EndoS on IgG glycosylation during streptococcal infections *in vivo*, we first needed an assay powerful enough to allow us to quantify the EndoS reaction products directly from very complex samples such as patient material (plasma, wound swabs or throat swabs). Previous attempts at measuring EndoS activity relied on SDS-PAGE^31,32^ or analysis of released glycans by HPLC^33^. These methods lack sensitivity, specificity and/or are ill suited for analysis of complex material. We therefore developed an SRM mass spectrometry method to specifically quantify the amount of glycan-hydrolyzed IgGs (IgG_GH_). As EndoS cleaves within the chitobiose core of the IgG *N*-glycan (Fig. 1a), it leaves the reducing end *N*-Acetylglucosamine (GlcNAc) residue attached to the protein^34^ and leads to the generation of a new IgG proteoform with a truncated glycan that is not detected in healthy individuals^35,36^. Samples were digested with trypsin and the peptides were treated with α-fucosidase to remove potential core fucosylations and end up with a single EndoS reaction product per IgG subclass. We defined SRM transitions for the tryptic peptides containing the glycosylation site of human IgG (IgG1: EEQYN(GlcNAc)STYR, IgG2: EEQFN(GlcNAc)STFR, IgG3: EEQYN(GlcNAc)STFR IgG4: EEQFN(GlcNAc)STYR) modified with a single *N*-linked GlcNAc residue. As the glycopeptides of IgG3 and IgG4 have the same precursor mass, we defined transitions that are in common and quantified both peptides together. To assess total IgG levels, previously developed SRM assays for quantification of each IgG subclass^37^ were included. All the transitions are listed in Table S2. To determine the absolute amounts of IgG and IgG_GH_, we synthesized heavy-isotope labeled standard peptides corresponding to the peptides we analyzed, which could be spiked into the samples to act as internal standards (Fig. 1b). The method was calibrated using a defined human standard serum (Fig. S1&2, Table S3) and validated by analyzing a set of human blood plasma samples with defined amounts of IgG_GH_. For method comparison, the same samples were also analyzed by SDS-PAGE and *Lens culinaris* agglutinin (LCA) lectin blot (Fig. 1cd). The SRM method exhibited better precision and accuracy as well as a much lower detection limit. Especially for samples where the IgG_GH_ content was low, as might be expected in clinical samples, the SRM method outperformed the SDS-PAGE assay. This together with low sample requirements, a large dynamic range and high analytical precision (Fig. S1, S2) made this method highly suitable for the analysis of complex patient materials. With this, method development was complete and we could start measuring EndoS activity during GAS infections *in vivo*.

### IGG GLYCANS ARE HYDROLYZED DURING GAS TONSILLITIS

Tonsillitis is the most common form of GAS infection and is characterized by throat pain, fever, tonsillar exudates and cervical lymph node adenopathy^1^. To study the effects of EndoS on patient IgGs during such an infection, we obtained 59 throat swab samples from a total of 54 patients who sought medical attention for a sore throat (Fig. 2a). 26 of these were diagnosed with GAS tonsillitis by rapid strep test and/or throat culture and were prescribed oral antibiotics. The other 28 patients exhibited a negative strep test and throat culture; therefore the infection was suspected to be viral and left untreated. 5 of the patients diagnosed with GAS tonsillitis were willing to return after antibiotic treatment and an additional throat swab was collected for each of these (Fig. 2a)

**FIGURE 2:**
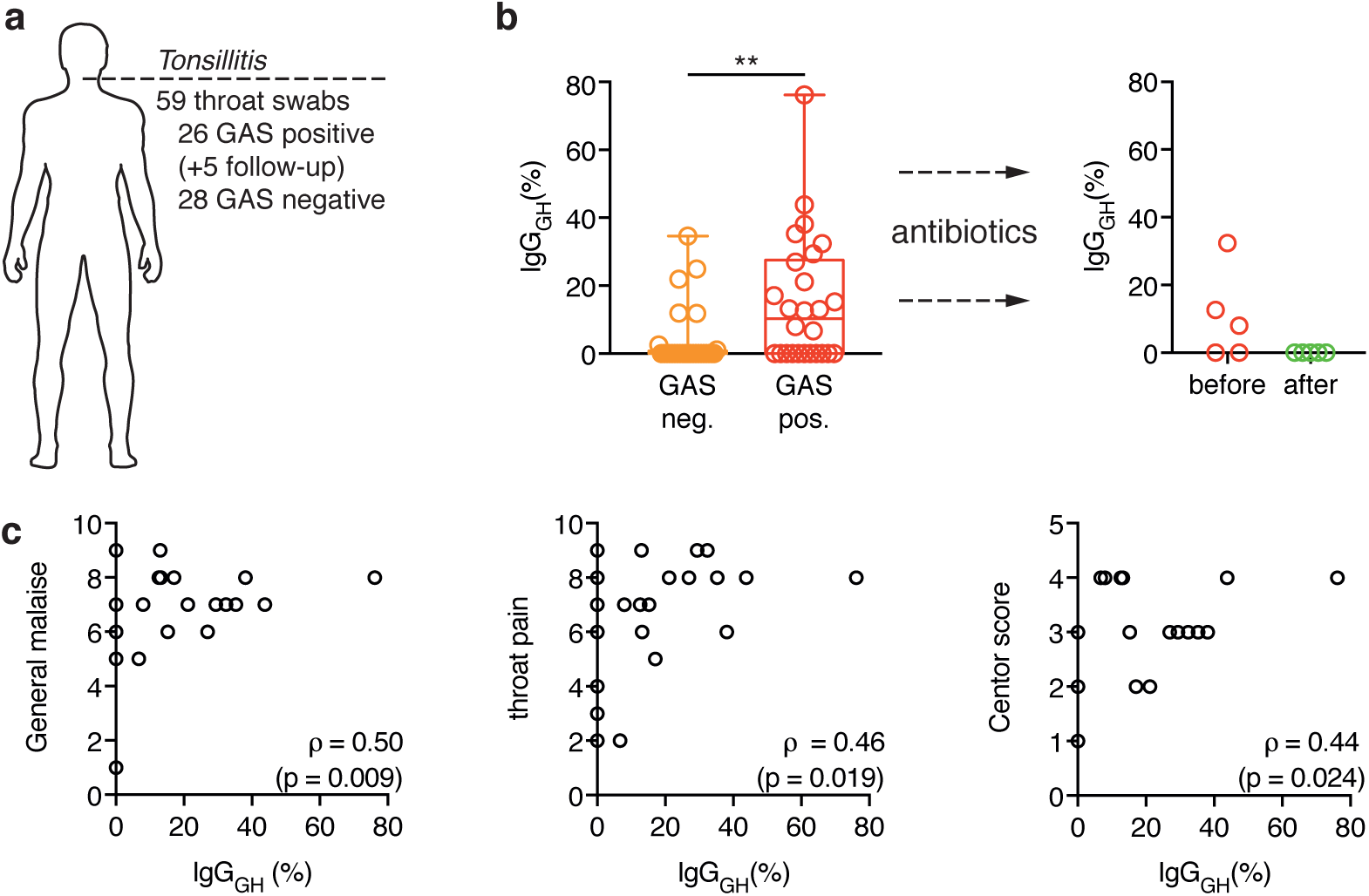
IGG GLYCAN HYDROLYSIS DURING GAS TONSILLITITS. (a) Overview of the collected throat swab samples from patients seeking medical attention for a sore throat. A total of 59 samples were taken from 54 different patients (26 GAS positive tonsillitis, 28 GAS negative tonsillitis). Follow-up refers to additional samples that were taken from 5 of the GAS tonsillitis patients after antibiotic treatment. (b) Percentage of glycan-hydrolyzed IgG as determined by SRM-MS analysis of tonsillar swabs from patients with, either GAS-negative (orange) or GAS-positive (red) tonsillitis. The boxes represent the 25^th^ to 75^th^ percentile with the median depicted as a line in the middle. The whiskers reach from the smallest to the largest data point, all of which are marked as circles. Glycan hydrolysis of the individual subclasses is shown in table S4. The glycopeptides from IgG3 and IgG4 could not be measured in these samples due to interfering background and were omitted from this analysis. Data was analyzed using a Mann-Whitney test (Table S6) (ns: p>0.05, **: p<0.01). (c) The tonsillitis patients were asked to grade their a general malaise (left) and throat pain (middle) on a scale from 0-10 and the centor score^38^ (right) was determined. These parameters were correlated to the IgG glycan hydrolysis measured in tonsillar swabs using SRM-MS. Correlation was analyzed according to Spearman (Table S7).

We used SRM mass spectrometry to quantitatively analyze the levels of IgGs as well as their glycosylation status in these throat swab samples. We observed no difference between GAS-positive and GAS-negative patients in total IgG content or in the distribution of the four IgG subclasses (data not shown). However, the percentage of glycan-hydrolyzed IgGs was significantly higher in the GAS tonsillitis group, with glycan hydrolysis approaching 80% in one case (Fig. 2b, left panel). This was no longer detectable in any of the samples taken after antibiotic treatment (Fig. 2b, right panel). Furthermore, IgG glycan hydrolysis could be correlated to the grade of throat pain and the general malaise experienced by the patients as well as their Centor scores (a clinical scoring system aimed at distinguishing GAS tonsillitis from viral infections^38^) (Fig. 2c). Taken together, these results suggest that EndoS is expressed and active during acute GAS tonsillitis, but its effects quickly disappear upon therapeutic intervention.

### IGG ANTIBODIES ARE DEGLYCOSYLATED SYSTEMICALLY DURING GAS INDUCED SEPSIS

Sepsis is a state of systemic inflammation in response to an infection and the worst case scenario in GAS infections^1^. To address the effect of EndoS on patient IgG antibodies during invasive GAS infection leading to sepsis, we collected blood plasma from 32 patients suffering from sepsis of various degrees of severity as well as 12 healthy volunteers. 18 patients had confirmed GAS infections whereas the other 14 suffered from various other bacterial infections (Fig. 3a, table S5). All the sepsis patients were ranked according to the degree of severity of their condition (sepsis, severe sepsis or septic shock). We used our SRM method to determine IgG levels and glycosylation state in blood plasma (Fig. 3b, table S5). The total IgG levels were not significantly different between the groups (Fig. 3b) and no appreciable amounts of glycan-hydrolyzed IgG could be detected in any of the plasma samples from the control groups (healthy or non-GAS sepsis). The same was true for milder cases of GAS induced sepsis. However, the plasma of 5 out of 6 GAS patients suffering from septic shock contained significantly increased amounts of IgG_GH_ (up to 1 mg/ml in the most severe case) (Fig. 3b).

**FIGURE 3:**
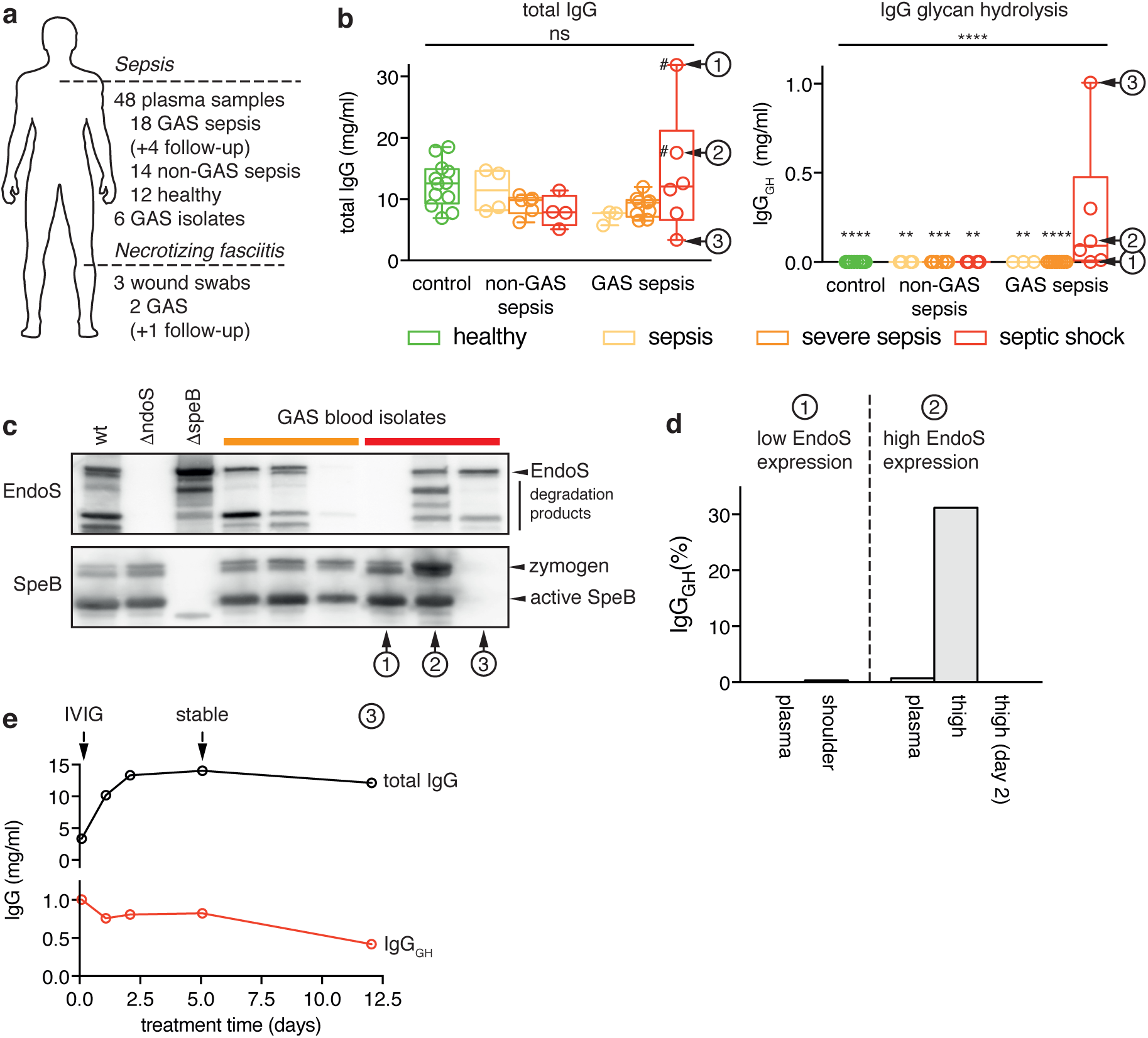
IGG GLYCAN HYDROLYSIS DURING INVASIVE GAS INFECTION. (a) Overview of the collected samples from sepsis patients. A total of 48 plasma samples, 3 wound swabs and 6 GAS isolates was collected from 32 patients (18 GAS sepsis, 14 non-GAS sepsis) and 12 healthy control individuals. Follow-up refers to 4 additional plasma samples that were taken from the same patient during the course of treatment and recovery. (b) Plasma concentration of total IgG (left panel) as well as the glycan-hydrolyzed IgG (right panel) as determined by SRM-MS. The patients are grouped according to infection state (healthy, non-GAS sepsis, GAS sepsis), as well as sub-grouped according to severity of disease (sepsis, severe sepsis, septic shock). The boxes represent the 25^th^ to 75^th^ percentile with the median depicted as a center line. The whiskers reach from the smallest to the largest data point, all of which are marked as circles. Glycan hydrolysis of the individual subclasses is shown in table S5. The p-value of the overall comparison of all the groups (by Kruskal-Wallis test) as well as adjusted p-values for the individual comparisons of the GAS septic shock group with each of the other groups are depicted (ns: p>0.05, *: p<0.05, **: p<0.01, ***: p<0.001, ****: p<0.0001). For a more detailed description of the statistical analysis see tables S8&S9). Two patients (marked by the hashtag #) had received IVIG treatment before the sample was drawn, affecting their total IgG concentrations. From three patients (marked 1-3) additional samples could be obtained and their analysis is shown in the other panels of this figure. (c) Expression of EndoS and SpeB by clinical GAS isolates *in vitro*. Blood culture isolates from patients 1-3 as well as three patients from the GAS severe sepsis group were analyzed with respect to their ability of secrete EndoS (top panel) and SpeB (bottom panel) into the culture supernatant *in vitro*. The culture supernatants were analyzed by SDS-PAGE followed by immunoblotting using rabbit antisera specific to EndoS and SpeB respectively. AP1 (wt) and isogenic *ndoS* and *speB* mutants were used as positive and negative controls respectively. (d) Local vs. systemic glycan hydrolysis. From patients 1 & 2, wound swab samples from the site of infection (patient 1: shoulder and patient 2: thigh) could be obtained and were analyzed by SRM-MS. The percentage of IgG_GH_ in the tissue as well as in plasma is shown. (e) IgG glycan hydrolysis over the course of infection. A series of plasma samples from patient 3 (starting at 2h after admission until 12 days later) was analyzed using SRM-MS. Both the concentration of total IgG (black) as well as IgG_GH_(red) is shown. The arrows mark onset of IVIG treatment (4h) and the time point when the patient was stable (5 days).

As we observed large differences in IgG glycan hydrolysis, even among the septic shock patients, we hypothesized that differential expression of EndoS among the different GAS strains infecting these patients could account for part of the observed variance. We were able to obtain 6 GAS isolates from the blood cultures of these patients and analyzed the ability of these strains to secrete EndoS into the culture medium *in vitro*. 3 isolates from severe sepsis patients and 3 from septic shock patients (Fig. 3c, patients 1, 2 and 3) could be obtained. The strains exhibited a large variability in EndoS expression as well as different levels of degradation of the EndoS protein. Strains secreting substantial amounts of EndoS and strains secreting almost no EndoS could be found in both groups. However, the amount of EndoS secreted by the isolates from the septic shock patients *in vitro* corresponded well with the amount of glycan-hydrolyzed IgG found in the corresponding patient’s blood plasma. The isolate from the patient with no detectable IgG glycan hydrolysis *in vivo* did not secrete detectable amounts of EndoS *in vitro*, and conversely, the isolate from the patient with the highest *in vivo* glycan hydrolysis secreted the most EndoS *in vitro*. The cysteine protease SpeB is a secreted GAS virulence factor and has been shown to cleave the EndoS enzyme^39^. Strikingly, SpeB is absent from the culture supernatant of the GAS isolate from patient 3 (with the highest degree of plasma IgG glycan hydrolysis, Fig. 3b) and consequently EndoS is largely intact.

The IgG glycan hydrolysis we observed in plasma reflects a systemic modification of IgG, which, due to the abundance of the antibody in circulation, necessitates large amounts of IgG to be deglycosylated to reach detectable levels. Locally, at site of infection, the effects of EndoS on the IgG pool are likely to be much more pronounced. To test this, we obtained wound swabs from the infected tissue taken during surgery from two of the sepsis patients suffering from necrotizing fasciitis (patients 1 & 2). We analyzed them by SRM mass spectrometry to determine the degree of IgG glycan hydrolysis and compared the results to those previously obtained from analysis of plasma samples (Fig. 3d). The samples originated from two of the patients whose GAS isolates we have analyzed for EndoS expression *in vitro* (Fig. 3c, patients 1 & 2). One isolate did not secrete any detectable amounts (Fig. 3c, patient 1) whereas the other one exhibited high EndoS expression (Fig. 3c, patient 2). Accordingly, two very different patterns of IgG glycan hydrolysis could be observed in these patients (Fig 3d). The first patient showed no detectable IgG glycan hydrolysis in plasma and only a minor amount in the wound swab sample. The second patient on the other hand exhibited moderate IgG glycan hydrolysis in plasma (~0.7 % hydrolyzed) and the amount of IgG_GH_ was considerably higher in the wound swab sample from the same day (~30% hydrolyzed). This was no longer detectable in a sample from the same site that was taken during a second surgery the day after.

As we observed that local IgG glycan hydrolysis was transient, we wanted to determine how long-lasting the EndoS-mediated perturbation of the systemic IgG pool was. To this end we obtained further plasma samples taken throughout the treatment and recovery periods (until 12 days after admission) from the patient exhibiting the highest amount of IgG_GH_ in plasma (Fig. 3b, patient 3). Shortly after admission (time point 2h), the patient presented with very low total IgG levels (3.3 mg/ml) and a high degree of IgG glycan hydrolysis (1 mg/ml). The patient was given intravenous immunoglobulin (IVIG) treatment, upon which the total IgG levels quickly normalized but the concentration of IgG_GH_ stayed high throughout the analyzed time interval and was still around 0.5 mg/ml at the 12 days end point (Fig. 3e).

Taken together, these results show that EndoS is expressed and active during acute GAS infection *in vivo*. It is able to hydrolyze the glycans from a considerable portion of the IgG pool locally at the site of infection (both in tonsillitis and necrotizing fasciitis) as well as systemically in the most severe cases of GAS sepsis. This points towards an important role for EndoS in evasion of the immune defenses by perturbation of the hosts IgG response.

### ENDOS IS EXPRESSED DURING GROWTH IN SALIVA AND PROTECTS GAS FROM PHAGOCYTIC KILLING

While we were able to show that substantial IgG glycan hydrolysis takes place during GAS infections, the functional consequences of this process for GAS pathogenesis remained unclear and needed to be studied using appropriate model systems. As EndoS activity was measureable in the majority of samples from GAS tonsillitis patients, we attempted to set up a simplified *in vitro* model reminiscent of the conditions GAS encounters on an inflamed tonsil during such an infection. When GAS colonizes the throat it would encounter saliva, an increasing amount of plasma proteins as inflammation leads to vascular leakage^37^ and finally phagocytic immune cells trying to eradicate the bacteria. To approximate these conditions, we grew GAS strains 5448 and AP1 as well as their respective isogenic *ndoS* mutants in sterile-filtered saliva supplemented with 5% serum and tested if EndoS would be expressed and active by SDS-PAGE. Based on electrophoretic mobility and/or LCA reactivity, both wild type strains but neither of the mutants were able to fully deglycosylate the IgG pool under these conditions (Fig. 4a, Fig. S6). Strain AP1 further expressed IdeS^11^ leading to proteolytic cleavage of IgG hinge in the culture supernatant. This resulted in a population of antibodies where the Fc glycans and at least one half of the heavy chains had been cleaved (Fig. S6).

**FIGURE 4:**
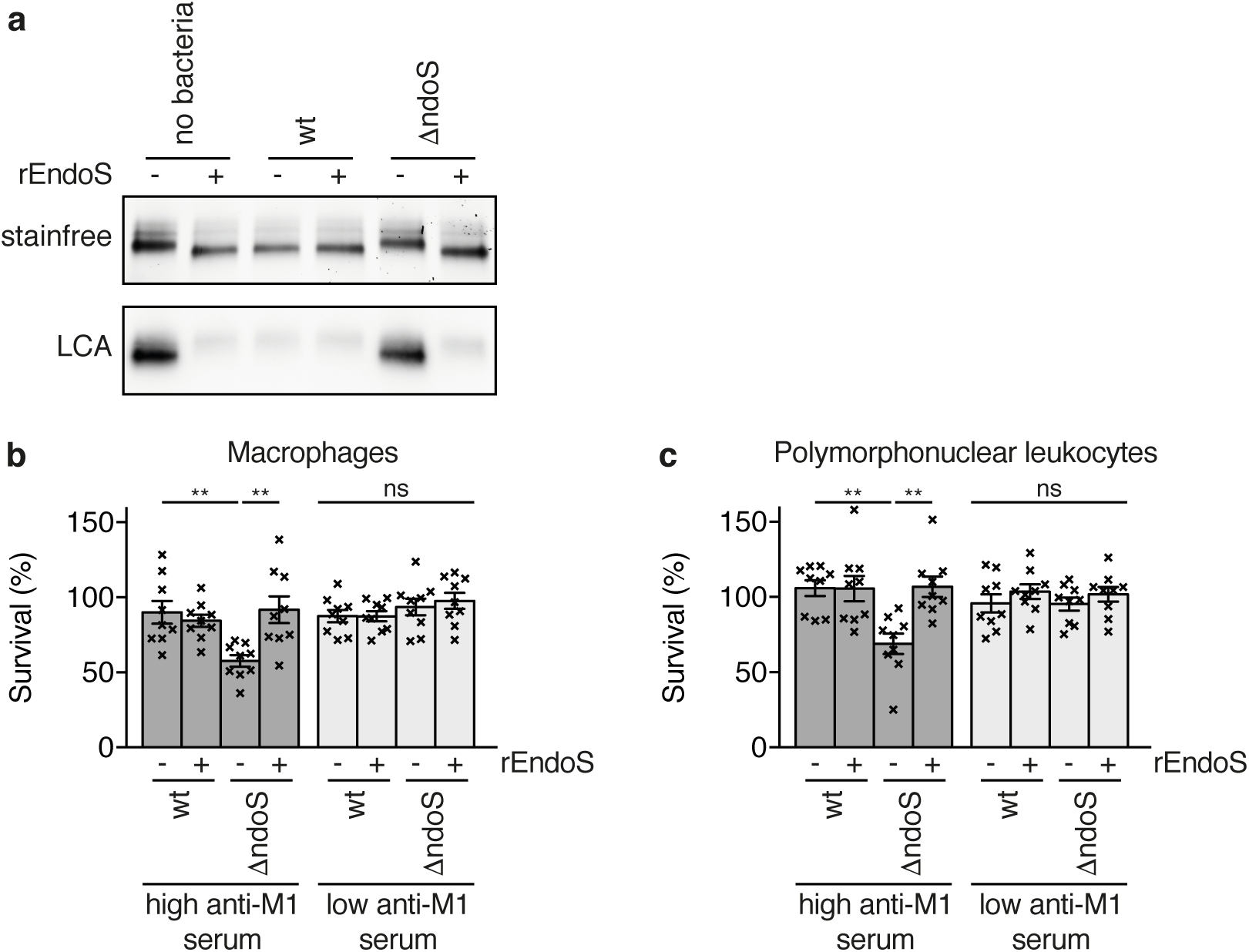
ENDOS CONFERS RESISTANCE TO PHAGOCYTIC KILLING. (a) GAS 5448 and an isogenic *ndoS* mutant were grown in human saliva supplemented with 5% human serum. IgGs were purified by Protein G and analyzed by SDS-PAGE (top) and LCA blot (bottom). Addition of recombinant EndoS (rEndoS) was used to complement the mutation. (b) Saliva-grown GAS 5448 and an isogenic *ndoS* mutant were challenged with human monocyte-derived macrophages (left) and human polymorphonuclear leukocytes (right) in the presence of serum with a high (dark grey) or low (light grey) anti-M1 IgG response. Survival rates were determined by numerating bacteria both in the initial inoculum as well as after incubation with the phagocytic cells. Data from 3 independent experiments with different cell donors (each preformed in triplicate, total n=9) was combined and analyzed using a Kruskal-Wallis test followed by Dunn’s multiple comparison test (Tables S10-13) (ns: p>0.05, **: p<0.01). The bar represents the mean, with the standard error depicted as error bars. Each individual data point is represented with a cross, showing the variability between the individual experiments and the replicates within the same experiment.

As IdeS and EndoS might be partially redundant and strain 5448 only secreted EndoS without any detectable IdeS activity when grown in saliva, we used this strain to test the resistance of wild type and *ndoS* mutant bacteria to phagocytic killing by human monocyte-derived macrophages (MDMs) and human polymorphonucelar leukocytes (PMNs) under the conditions described above. Deletion of the *ndoS* gene led to a small but significant increase in killing of the bacteria by both MDMs and PMNs (Fig. 4bc) and addition of recombinant EndoS reversed the phenotype. However, increased susceptibility to phagocytic killing was only observed when the assay was performed in the presence of serum containing GAS-specific IgGs (as determined by measuring the IgG response to streptococcal M1 protein by ELISA (Fig. S5)). In the absence of specific IgGs both mutant and wild type exhibited similar resistance to phagocytic killing. This indicates that EndoS confers increased resistance by neutralizing specific IgGs directed towards the pathogen and prevents them from mediating phagocytosis.

### ENDOS MUTANT GAS ARE LESS VIRULENT IN A MOUSE MODEL OF INVASIVE GAS INFECTION

In order to study the role of EndoS in neutralization of GAS-specific IgGs in more detail and determine its contribution to the outcome of streptococcal infection, we established a mouse model. As IdeS has no discernible activity on relevant subclasses of murine IgG (IgG1 and IgG2b)^40^ we were able to use the more mouse-virulent strain AP1 for these experiments without confounding the results. Wild type and *ndoS* mutant^34^ GAS were used to infect C57BL/6J mice subcutaneously and both local (skin) and systemic (plasma, spleen) samples were taken at 48h post infection to determine bacterial loads as well as IgG glycan hydrolysis by SRM mass spectrometry (Fig. 5a). An assay analogous to the one for human IgGs was developed to quantify murine IgG1 levels and its glycosylation status (Fig. S5). As EndoS does not exhibit any murine IgG subclass specificity^41^, this can be used as an indicator for overall IgG glycan hydrolysis. When mice were infected with a wild type GAS strain, IgG1 was almost completely deglycosylated locally and around 30% glycan hydrolysis was observed systemically (Fig. 5b, S8). The animals exhibited a heterogeneous response to infection, with greatly varying degrees of severity observed. Consequently the measured levels of IgG glycan hydrolysis also showed a large variance. We therefore tried to correlate IgG glycan hydrolysis and bacterial load in the skin samples and found almost perfect correlation (Fig. S5). Mice infected with the *ndoS* mutant developed local and systemic signs of infection to a similar degree but showed no detectable IgG glycan hydrolysis (Fig. 5b, S7,8). This indicates that EndoS is expressed and active during such an infection but does not confer any selective advantages under these naïve conditions.

**FIGURE 5:**
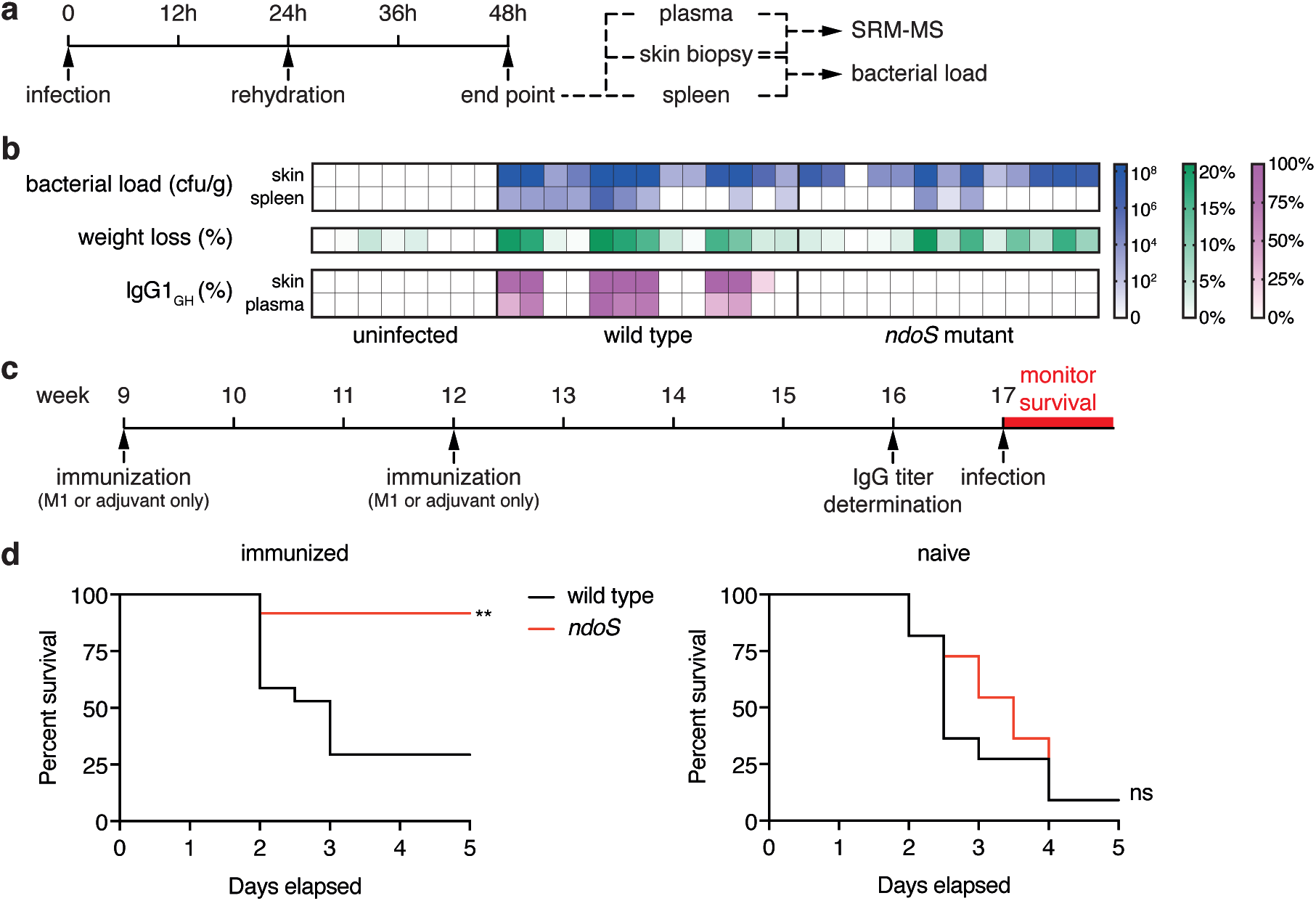
ENDOS LEADS TO IGG GLCYANS HYDROLYSIS AND CONFERS A SELECTIVE ADVANTAGE IN A MOUSE MODEL OF GAS INFECTION. a) Experimental setup for the infections of immunologically naïve mice. 9 week old, female C57BL/6 mice were infected subcutaneously in the flank with GAS AP1, an *ndoS* mutant or PBS. The animals were rehydrated by injection of 0.5 ml saline at 24 h and sacrificed at 48 h post infection. The spleen and a skin biopsy from the site of injection were taken to assess bacterial loads and the skin sample, as well as a plasma sample were used for analysis by SRM-MS. b) Bacterial load (blue), weight loss (green) and percentage of IgG1 glycan hydrolysis as determined by SRM-MS (purple) at 48h post infection. Each column corresponds to an individual animal. c) Experimental setup for the immunization with M1 and subsequent infection of mice. 9 week old, female C57BL/6 mice were injected with purified M1 protein or adjuvant only and received a second dose at 12 weeks of age. After 4 weeks, the anti-M1 IgG response was assessed by ELISA and the animals were infected subcutaneously one week later. d) Survival of M1-immunized (right) and mock-immunized (left) animals after infection with either GAS AP1 wild-type (black) or *ndoS* mutant (red). Mice were monitored twice daily for survival for a period of 5 days. Curves were compared using a Mantel-Cox test (ns: p>0.05, **: p<0.01).

As EndoS targets the adaptive immune response, its functional role during GAS infection is most appropriately studied in the context of adaptive immunity. We therefore immunized mice prior to infection through two injections with purified streptococcal M1 protein combined with adjuvant. After an IgG response to M1 was confirmed, the mice were infected subcutaneously and survival was monitored for 5 days (Fig. 5c). This immunization protocol lead to a complete protection at an infectious dose of 2.5×10^5^ cfu which could be overcome by increasing the dose to 2×10^7^ cfu (Fig. S9). The majority of mice infected with wild type bacteria at that dose succumbed to the infection within 2-3 days. On the other hand, more than 90% of the mice infected with *ndoS* mutant bacteria survived the infection with only milder symptoms (Fig 5d, left). This profound difference in susceptibility to infection between wild-type and mutant could not be observed in animals that were mock immunized by injection of adjuvant only, where both groups showed a similarly high mortality (>90%, Fig. 5d, right).

## Discussion

IgG is a central molecule of the mammalian immune system. It provides a link between adaptive and innate immunity by specifically binding to antigens presented by a pathogen with its Fab regions and recruiting immune effectors with its Fc region. Which exact effector functions are elicited is a highly regulated process that lets the immune system tune its response to the pathogen in question and mediate different effector functions against for example a gram-positive bacterium, a gram-negative bacterium or a virus. One determining factor in this process is the nature of the IgG antibody itself; namely its subclass and the structure of its Fc glycosylation.

We here present the first evidence of a pathogen exploiting this regulatory mechanism by specifically altering IgG Fc glycosylation *in vivo* during the clinical course of an infection. By implementing a targeted proteomics approach based on SRM-MS we were able to study the effects of EndoS on IgG antibodies during natural infections in human patients. Using SRM we were able to detect and absolutely quantify glycan-hydrolyzed IgGs in a subclass specific manner directly from a variety of very complex patient samples. With only minimal amounts of sample required and detection limits below 0.5 ng, this method is far superior to any other techniques used to measure IgG glycan hydrolysis to date. With this, we were able to quantitatively address the effects of EndoS on a patient IgG population during streptococcal infections. While a slight preference of EndoS towards IgG1^42^ has been reported, we did not observe any subclass specificity in any of the patient samples (Table S4,5). Locally, at the site of infection, the degree of IgG glycan hydrolysis was generally high (up to nearly complete hydrolysis) but the effects were only transient, disappearing quickly upon therapeutic intervention. This was true both for mild infections (tonsillitis) as well as severe, invasive infections (necrotizing fasciitis). This indicates that at site of infection, IgG is turned over quickly and constantly replenished from the circulation and IgG glycan hydrolysis is only detectable as long as bacterial load is high and EndoS is continuously secreted. Due to the much larger amount of IgG in circulation as compared to the infected tissue and the increased dilution of the enzyme further away from the site of infection, effects on the systemic IgG pool in circulation were much harder to detect and therefore rarer. Only in the most severe cases of invasive infections (septic shock), could glycan hydrolysis be observed systemically. This systemic IgG glycan hydrolysis was long lasting with approximately half of the glycan-hydrolyzed IgGs still present after 12 days. A similarly slow recovery of IgG glycosylation to normal levels was also observed when rabbits or mice were injected with recombinant EndoS as an experimental treatment of autoimmunity^31,43^.

In both of our patient sets, only a portion of the patient samples showed detectable levels of IgG_GH_ (58% of tonsillitis patients locally and 28% of sepsis patients systemically). In part, this might be due to some GAS isolates expressing only very low levels of EndoS (Fig. 3c). *In vivo* glycan hydrolysis was not associated with *covRS* mutation or any specific *emm*-type. Indeed we detected it in infections with at least different 8 different *emm*-types (Table S4,5). All publicly available GAS genome sequences contain an *ndoS* or an *ndoS*-like gene^44^, but expression of the EndoS protein differs greatly between different GAS isolates. However, the fact that IgG glycan hydrolysis correlates to disease severity both in tonsillitis and invasive disease indicates that we – due to analytical limitations – might not be able to detect EndoS-mediated IgG glycan hydrolysis in samples from patients suffering from less severe infections. This is true especially for the GAS sepsis patient samples. Apart from two exceptions, we were only able to study IgG in blood plasma and therefore were unable to address potential perturbations of the local IgG pool at site of infection in more detail. Indeed among the sickest patients in each cohort the percentage of samples with detectable IgG_GH_ was considerably higher. 5 out of 6 GAS septic shock patients (83%) exhibited measureable systemic IgG glycan hydrolysis and among the sickest half of the tonsillitis patients (based on estimation of general malaise) glycan hydrolysis of the local IgG pool was detectable in all but one patient (92% total). While studying the activity of EndoS during GAS tonsillitis, we also observed a considerable amount of IgG glycan hydrolysis in some of our control samples. We speculate that this was due to enzymatic activities of the oral micro flora, especially oral streptococci which are known to express a large number of glycoside hydrolases^45^. This is supported by the fact that IgG glycan hydrolysis was not detectable in any of the samples we took after antibiotic treatment.

While we were able to show that IgG glycan hydrolysis takes place during GAS infection *in vivo*, the functional consequences of this process could not be deduced directly from the patient data. Removal of the Fc glycan by EndoS has previously been shown to impair both Fc receptor interaction and complement activation *in vitro*^20,25^. While EndoS has been speculated to contribute to GAS virulence, all studies showing this to date had to resort either to addition of recombinant EndoS or overexpression^25,26^ to see any effect. Thus, the conditions under which endogenous EndoS is expressed, active and able to confer a selective advantage to the bacteria remained unclear. EndoS expression is highly regulated and while the regulatory network responsible remains obscured, it has been shown to involve both transcriptional regulation by *ccpA* and weakly by *covR/S*^46^ as well as posttranslational regulation through proteolysis^47^. Transcriptomics studies have shown that EndoS expression is repressed during growth in rich medium^48^ (such as the standard Todd-Hewitt broth) and is not induced until the bacteria reach stationary phase^49^. This means that most standard assays used to study streptococcal virulence factors such as incubating log phase bacteria with phagocytic cells and IgG (or the classic Lancefield assay^50^) are ill suited to address the role of EndoS or similar secreted immunomodulatory activities. This might constitute a major shortcoming in for example analysis of protective effects of IgGs in vaccination studies, potentially resulting in overestimation of vaccine efficacy.

We used our characterization of IgG glycan hydrolysis *in vivo* to set up relevant model systems to address the contribution of EndoS to GAS pathogenesis. Based on our findings on IgG glycan hydrolysis during GAS tonsillitis, we were able to approximate the conditions GAS encounters on an inflamed tonsil using human saliva and serum. This prompted wild type bacteria to secrete enough EndoS to completely deglycosylate the IgGs present and neutralize the contribution of GAS-specific IgGs to phagocytic killing by human macrophages or neutrophils *in vitro*. *ndoS* mutant bacteria on the other hand were unable to cleave the glycans from IgG and consequently exhibited a higher susceptibility to phagocytic killing. This phenotype could be reversed by externally adding recombinant EndoS. These results, together with the fact that the *ndoS* mutation had no phenotype in the absence of GAS-specific IgGs indicates that the observed phenotype is due to deglycosylation of the GAS-specific IgGs which in turn impairs their ability to mediate phagocytic killing. There seems to be no general attenuation of the *ndoS* mutant, nor any benefits in glycan hydrolysis of non-specific IgGs.

An animal model of local skin infection leading to invasive infection showed very similar dynamics. Mice infected with *ndoS* mutant bacteria exhibited no IgG glycan hydrolysis and were significantly less likely to die from the infection than mice infected with wild type GAS. This difference was however only clearly evident in the context of adaptive immunity (i.e. in animals that had been immunized against GAS prior to infection). In agreement with the results from the phagocytosis assays and previous studies^26^, naïve mice showed a very similar susceptibility towards both wild type and *ndoS* mutant bacteria; with no significant differences in survival, weight loss or bacterial load in the skin (Fig. 5, Fig. S7). Only the bacterial burden in the spleen was slightly decreased in mice infected with *ndoS* mutant bacteria (Fig. S7). This might point to a small degree of innate protection conferred by natural IgGs^51^ that can be counteracted by EndoS.

While not generally used to treat sepsis^52,53^, administration of intravenous immunoglobulin (IVIG) has shown promise as a treatment for streptococcal toxic shock syndrome^54,55^ and necrotizing fasciitis^56^. Its mode of action is thought to involve neutralization of bacterial superantigens and inhibition of pro-inflammatory signaling^57–59^. Our data points to another reason why IVIG treatment could improve the outcome of severe streptococcal infections. EndoS-mediated glycan hydrolysis inactivates IgGs and as shown in this study, such antibody modifications can be systemic, long lasting and affect a considerable fraction of the patient’s total IgG pool. GAS can lower functional IgG levels even further through secretion of the IgG protease IdeS^11,37^. In such cases, IVIG in concert with antimicrobial therapy could help to quickly normalize the level of functional IgG in circulation. We observed such an effect in one patient (Fig. 3e, patient 3).

Despite an abundance of anti-GAS IgGs in the serum of most people^4–7^, protective immunity towards the pathogen does not ensue. Furthermore, development of effective GAS vaccines has proven challenging. Herein we present a possible mechanism to explain why anti-GAS IgGs often confer such poor protection: GAS is able to effectively neutralize the contribution of IgG to host defense through specific degradation of the IgG antibodies. We have demonstrated the dynamics of IgG glycan hydrolysis by EndoS in this study and a recent study has also shown significant levels of IdeS-mediated IgG proteolysis during GAS infections *in vivo*^37^. This makes GAS a very proficient evader of IgG-mediated immunity, a fact that has implications for the treatment of severe GAS infections and has to be taken into account in future research into the immune response to GAS as well as in the continuing efforts at development of an effective vaccine against the pathogen. Finally, immune evasion through modification of IgG glycosylation might not be restricted to GAS and enzymes similar to EndoS can be found in many other pathogenic species such as *Enterococcus faecali*s^60^, *Streptococcus pneumoniae*^61^ as well as other non-group A streptococci^62,63^. Indeed, an EndoS homolog from *Streptococcus equi*^63^ proved a protective antigen in a vaccination trial in mice, pointing towards its importance for the infection process. Bacterial modulation of IgG glycosylation might therefore be a more widespread phenomenon that warrants further study.

## Materials and Methods

### PATIENT SAMPLES

Tonsillar swabs (n = 54, ESwab Liquid Amies) were obtained from patients (>8 years old) seeking clinical care because of a sore throat at the primary healthcare clinics at Laurentiikliniken and Skåne University Hospital (SUS) both in Lund, Sweden. GAS tonsillitis was diagnosed by rapid strep test (antigen detection) and routine bacterial culturing. A follow-up tonsillar swab sample (3-5 days post) was taken from a subset of patients (n=5) treated with antibiotics (Table S4). Wound swabs from the local infection site of patients clinically diagnosed with GAS sepsis and necrotizing fasciitis were obtained from SUS, Lund Sweden (n=4) during surgical intensive care (Table S5). Patient swab samples were transported on dry ice before storage at −80 °C. The patient plasma samples were part of a larger cohort collected at the Clinic for Infectious Diseases or at Intensive Care Unit at Lund University Hospital between 2005 and 2015. All human samples were obtained with informed consent and with the approval of the local ethics committee (see ‘Ethical Considerations’ section below).

### SAMPLE PREPARATION FOR MASS SPECTROMETRY

Proteins from the swab samples were extracted and homogenized in water using a bead-beater (Fastprep-96, MP-Biomedicals). 0.625 µl plasma or 50 µg of protein from swabs or skin homogenates were prepared for MS analysis using the SmartDigest Kit (Thermo Scientific). Samples were denatured at 90°C followed by digestion for 3.5 hours at 70°C. The peptides were reduced using 50 mM TCEP, alkylated with 100 mM iodoacetamide and finally purified using SOLAµ HRP plates (Thermo Scientific). The peptide samples were dried in a vacuum centrifuge and dissolved in 100 µl 50 mM sodium acetate buffer (pH 5) containing 50 mU Thermatoga maritima α-fucosidase (Megazymes). After incubation at 70°C for 14 h, the samples were purified a second time on SOLAµ HRP plates and dried in a vacuum centrifuge.

### SRM MASS SPECTROMETRY

Peptide samples were dissolved in 2%ACN, 0.2% FA and AQUA peptides (Thermo HeavyPeptide QuantPro, table S3) were spiked in. The amount of peptide standards was adjusted so that the ratio of light to heavy signal falls within the interval of 0.1 to 10. Sample corresponding to 1 µg of protein were analyzed by SRM mass spectrometry using a TSQ Vantage triple quadrupole mass spectrometer coupled to an Easy-nLC II system (both Thermo Scientific) equipped with a PicoChip column (PCH7515-105H354-FS25, New Objective). Data was acquired with a spray voltage of 1500V, 0.7 FWHM on both quadrupoles and a dwell time of 10ms. Assays for all the non-glycopeptides were obtained from published studies ^37,64^ while glycopeptide assays were developed as described here^65^. Assay set-up, empirical collision energy optimization as well as data analysis was done using Skyline ^66,67^. The analyzed transitions are listed in table S2.

### DETERMINATION OF DETECTION LIMITS

Human IgG subclass reference serum NOR-1 (NordicMUBio) was used to create standard samples for method calibration and determination of detection limits. For a fully glycan hydrolyzed sample, the serum was incubated with 50 µg/ml recombinant EndoS at 37°C for 16h. Both treated and untreated serum samples were prepared for mass spectrometry analysis separately as described above. Peptides were dissolved in 2%ACN, 0.2% FA at a concentration of 1 µg/µl and spiked with AQUA IgG glycopeptides. The EndoS-treated sample was further spiked with AQUA IgG peptides. Three separate dilution series were prepared by serially diluting the EndoS-treated sample with peptides from the untreated sample. Finally, peptides samples corresponding to 1 µg of protein form each dilution were analyzed by SRM mass spectrometry. From this data set detection limits for each peptide were determined separately:

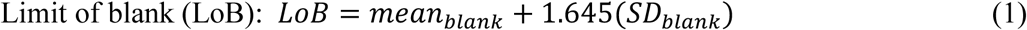

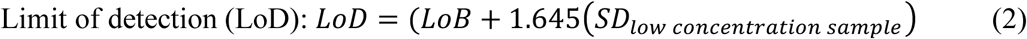

Untreated samples were used to determine LoB according to formula 1. The first dilution with a concentration above LoB was used as the low concentration sample for LoD determination with formula 2. A lower limit of quantification (LLoQ) was defined as two time LoD and in all further experiments, measured amounts below LLoQ were considered as being 0. As the concentrations of all 4 IgG subclasses in serum NOR-1 have been determined by the manufacturer, we could also use this dataset to calibrate the quantification and correct for incomplete digestion and sample loss during preparation. For each data point above LOD and with a light to heavy between 0.1 and 10 conversion factors were determined by dividing the known IgG subclass concentrations with the measured light to heavy ratios. These conversion factors were averaged (table S3) and are used to determine absolute amounts from measured light to heavy ratios in all further experiments with human samples. The same procedure was followed when the assay for murine IgG1 was developed. The standard sample there was a monoclonal mouse IgG1 (MA-69, Biolegend, San Diego, USA) that was treated with recombinant EndoS and spiked into human plasma as a background proteome.

### SDS-PAGE AND LECTIN BLOT

IgG was purified using Protein G (Ab SpinTrap, GE Healthcare). Samples were separated on SDS-PAGE (Mini-protean TGX stainfree gels, 4-15% acrylamide, BioRad), imaged using a ChemiDoc MP imaging system (BioRad) and transferred to low fluorescent PVDF membranes using the Transblot Turbo kit (BioRad). Membranes were blocked for 1h in lectin buffer (10 mM HEPES pH7.5, 150 mM NaCl, 0.01 mM MnCl_2_, 0.1 mM CaCl_2_, 0.1% (v/v) tween 20) followed by incubation with 5 µg/ml fluorescein-labeled LCA lectin (Vector laboratories). After extensive washing in lectin buffer, the membranes were imaged using a ChemiDoc MP imaging system (BioRad).

### ANALYSIS OF ENDOS AND SPEB EXPRESSION *IN VITRO*

AP1, AP1ΔndoS, AP1ΔspeB and the GAS clinical isolates were grown overnight at 37°C, 5% CO_2_ in C-medium (0.5% (w/v) Proteose Peptone No. 2 (Difco) and 1.5% (w/v) yeast extract (Oxoid) dissolved in CM buffer (10 mM K_2_PO_4_, 0.4 mM MgSO_4_, 17 mM NaCl pH 7.5). The cultures were pelleted and the supernatants sterile filtered (Millex-GP filter unit 0.22μm, Millipore). Proteins were precipitated from the supernatants with 5% TCA (trichloroacetic acid) and analyzed by SDS-PAGE under reducing conditions. Proteins were transferred to PVDF-membranes using the Trans-Blot Turbo equipment (Bio-Rad Laboratories) according to manufacturer’s instructions. Membranes were blocked with 5% (w/v) blotting-grade blocker (Bio-Rad Laboratories) in PBST, followed by incubation with EndoS or SpeB rabbit antiserum ^12,68^ another wash and incubation with a secondary antibody (goat anti-rabbit HRP-conjugated antibody, BioRad). The membranes were developed using Clarity Western ECL substrate (BioRad) and visualized with a ChemiDoc MP Imager (Bio-Rad, USA).

### ANALYSIS OF *EMM* AND *COVRS* SEQUENCES

*emm* sequences of the GAS clinical isolates were analyzed according to protocols published by the CDC and compared to a database of known *emm* sequences using the tool on the CDC website (https://www2a.cdc.gov/ncidod/biotech/strepblast.asp). The *covRS* operon was sequenced as previously described^69^ and compared to published sequences of the same serotype.

### *IN VITRO* TONSILLITIS MODEL

#### Preparation of saliva

Saliva from healthy volunteers was collected in the morning after extensive brushing of the teeth. The saliva was centrifuged (20 min, 20000 g), sterile filtered (Steriflip GP 0.22 µm, Milipore) and either used directly or kept at −20°C until use.

#### Preparation of polymorphonuclear leukocytes (PMNs)

20 ml blood was collected into EDTA blood collection tubes (BD Bioscience) and the PMNs were isolated using PolyMorphPrep (Axis-shield) according to manufacturers recommendations. After counting, the cells were diluted into RPMI medium and seeded at 50000 cells/well into a 96-well plate.

#### Preparation of monocyte derived macrophages (MDMs)

Peripheral blood mononuclear cells (PBMCs) were isolated from leukocytes of healthy anonymous donors provided by the Lund University Hospital. Red blood cells were removed by centrifugation on Lymphoprep (Fresenius Norge As, Oslo, Norway) and recovered PBMCs were washed to remove platelets. Monocytes were isolated using a magnetic cell separation system with anti-CD14 mAb-coated microbeads (Miltenyi Biotec). CD14-positive monocytes were seeded into 12 well plates at 5 × 10^5^ cells/well and differentiated into macrophages by culture in complete RPMI 1640 medium (Gibco) supplemented with 10% heat-inactivated human AB+ serum, 50 nM β-mercaptoethanol (GIBCO), Penicillin-Streptomycin (Sigma) and 40 ng/ml M-CSF (Peprotech) at 37°C under a humidified 5% CO2 atmosphere for 6 days. Medium was replaced on day 2 and on day 4, when cells were washed with PBS and the medium was replaced with antibiotic-fee medium. The cells were further incubated until day 6 when the infection experiments took place.

#### Killing assays

GAS 5448 and an isogenic *ndoS* mutant^26^ were grown overnight in THY medium at 37°C, 5% CO_2_, diluted 1:10 into fresh medium and let grow to mid-log phase (OD = 0.4). Cultures were diluted 1:50 into 1 ml of saliva, incubated for 2h at 37°C, 5% CO_2_ and diluted again (1:20) into 1 ml fresh saliva (supplemented with 5% serum and 0.5 µg/ml recombinant EndoS where suitable). After 20h at 37°C, 5% CO_2_ the bacteria were diluted 1:10 into RPMI medium and used to infect PMNs or MDMs at an MOI of 2. After 30min (for PMNs) or 2 h (for MDMs) incubation at 37°C, 5% CO_2_ the cells were lysed using ddH2O (for PMNs) or 0.025% Triton X-100 (for MDMs) and the number of surviving bacteria was determined by plating on THY agar plates.

### ANALYSIS OF ENDOS EXPRESSION *IN VITRO*

AP1, the *ndoS* mutant and the GAS clinical isolates were grown in C-medium (37°C and 5% CO2) overnight and normalized to the same OD_620_ using fresh C-medium. Bacteria were pelleted by centrifugation and the supernatants filtered (Millex-GP filter unit 0.22μm, Millipore). Proteins were precipitated from the supernatants with 5% TCA (trichloroacetic acid) and analyzed by SDS-PAGE under reducing conditions. Proteins were transferred to PVDF-membranes using the Trans-Blot Turbo kit (Bio-Rad Laboratories) according to manufacturer’s instructions. Membranes were blocked with 5% (w/v) blotting-grade blocker (Bio-Rad Laboratories) in PBST, followed by incubation with EndoS^34^ or SpeB^68^ antiserum The membranes were washed followed by incubation with a secondary antibody (goat anti-rabbit HRP-conjugated antibody, BioRad). The membranes were developed using Clarity Western ECL substrate (BioRad) and visualized with a ChemiDoc MP Imager (Bio-Rad, USA).

### MOUSE INFECTIONS

*Acute infection model of GAS in naïve C57BL/6J mice* GAS AP1 and *ndoS* mutant (Table S1) were grown to logarithmic phase in Todd–Hewitt broth (37°C, 5% CO_2_). Bacteria were washed and resuspended in sterile PBS. 2-3×10^5^ cfu of AP1 (n=13) or *ndoS* mutant (n=13) were injected subcutaneously into the flank of 9-week-old female C57BL/6J mice (Scanbur/ Charles River Laboratories). Control mice were injected with PBS (n=8). Mice were rehydrated subcutaneously with saline at 24 h post infection. Body weight and general symptoms of infection were monitored regularly. Mice were sacrificed at 48 h post infection and organs (blood, spleens and skin) were harvested to determine the degree of bacterial dissemination and IgG glycan hydrolysis.

*M1 immunization and survival study* 9-week-old female C57BL/6J mice (Scanbur/ Charles River Laboratories) were injected subcutaneously with M1 protein on days 0 and 21 (10 μg/dose), purified as previously described^68,71^. The protein was administered as a 50:50 solution of M1:adjuvant (TiterMax Gold), in a 50 μl volume. Control mice were similarly injected with PBS:adjuvant solution. Serum was collected at day 49 and anti-M1 titers were measured by ELISA as previously described^72^ with a goat anti-mouse HRP-conjugated secondary antibody at 1:5000 (BioRad). Immunized and control mice were infected subcutaneously into the flank with 2×10^7^ cfu of the AP1 (n=17) or *ndoS* mutant (n=12) at day 56. Mice were rehydrated subcutaneously with saline 24 h post infection. Weight and general symptoms of infection were monitored every 12 h during the acute phase of the infection, and then every 24 h until termination of the study at 5 days post infection. Animals displaying a weight loss exceeding 20% (until 72 h post infection) or 15% (after 72 h post infection) were considered moribund and sacrificed.

### STATISTICS

All statistical analyses were preformed using GraphPad Prism 7. Phagocytic killing assays were analyzed using one-way ANOVA followed by Tukey’s multiple comparison test. IgG hydrolysis data was analyzed using a Mann-Whitney test or a Kruskal-Wallis test in combination with Dunn’s multiple comparison test (when more than two datasets were compared). Correlation was determined according to Spearman and survival data was analyzed using a Mantle-Cox test.

### ETHICAL CONSIDERATIONS

All animal use and procedures were approved by the local Malmö/Lund Institutional Animal Care and Use Committee, ethical permit number M115-13. Collection and analysis of human throat swabs and plasma samples was approved by the local ethics committee (dnr 2005/790, 2015/314 and 2016/39).

## AUTHOR CONTRIBUTIONS

AN designed the study, performed all mass spectrometry and phagocytosis experiments and wrote the manuscript. EB and OS designed and performed all animal experiments. CK helped with SRM assay development and data analysis. AL organized and supervised collection of clinical samples. RK prepared cells for phagocytosis experiments. JM and MC designed the study, supervised the research, and wrote the manuscript.

## DATA AVAILABILITY

The MS analysis files from Skyline underlying figures 1d, 2b, 3bde and 5b are available on PanoramaWeb (https://panoramaweb.org/endos.url). The MS raw data is available upon request.

### ACKNOWLEDGEMENTS

We thank Fanny Olsson Byrlind and Tomas Lindgren for help with collection of patient samples, Bo Nilsson for collecting the GAS clinical isolates, and Fredric Carlsson for helpful discussions about the choice of animal model. This work was supported by grants to AN from the Swiss National Science Foundation (P2EZP3_155594 and P300PA_167754), the Royal Physiographic Society in Lund and the Sigurd and Elsa Goljes Memorial Foundation. This work was further supported by grants to MC from the Swedish Research Council (projects 2012-1875 and 2017-02147), the Royal Physiographic Society in Lund, the Foundations of Åke Wiberg, Alfred Österlund, Gyllenstierna-Krapperup, Torsten Söderberg, the King Gustaf V`s 80 years fund, and Hansa Medical AB as well as grants to JM from Foundation of Knut and Alice Wallenberg (2016.0023), European research council starting grant (ERC-2012-StG-309831), the Swedish Research Council (project 2015-02481), the Wallenberg Academy Fellow program KAW (2012.0178 and 2017.0271), Olle Engkvist Byggmästare and the Medical Faculty of Lund University. The funders had no role in preparation of the manuscript or in the decision to publish

