## Supplemental figures, tables and methods for "Sweet revenge - *Streptococcus pyogenes* showcases first example of immune evasion through specific IgG glycan hydrolysis"

TABLE S1: STRAINS

| Strain | Description | Source |
|---|---|---|
| AP1 | *S. pyogenes* serotype M1, CovSΔD319-S500 | Collection of the WHO Collaborating Center for Reference and Research on Streptococci (Prague, Czech Republic) |
| AP1ΔndoS | Strain AP1 lacking the *ndoS* gene | Collin et al. 2001[1] |
| AP1ΔspeB | Strain AP1 lacking the *speB* gene | Collin et al. 2001[1] |
| 5448 | *S. pyogenes* serotype M1 originally isolated from a patient with necrotizing fasciitis, *covRS* wild-type | Chatellier et al. 2000[2] |
| 5448ΔndoS | Strain 5448 lacking the *ndoS* gene | Sjögren et al. 2011[3] |
| MC25 | Strain AP1 lacking the cell wall anchor of M1 protein, secretes M1 to the medium | Collin et al. 2000[4] |

### Table S2: SRM assay for human IgGs

Overview of all the SRM transitions analyzed in this study. All cysteines are carbamidomethylated. Underlined residues are heavy isotope-labeled ($^{13}C$ and $^{15}N$). The asterisk denotes fragment ions that have undergone a neutral loss of the GlcNAc modification.

| Protein | Peptide sequence | isotope label | precursor m/z | fragment m/z | collision energy | ion |
|---|---|---|---|---|---|---|
| human IgG1 | GPSVFPLAPSSK | light | 593.83 | 846.47 | 26 | y8 |
| | GPSVFPLAPSSK | light | 593.83 | 699.4 | 18 | y7 |
| | GPSVFPLAPSSK | light | 593.83 | 489.27 | 20 | y5 |
| | GPSVFPLAPSSK | light | 593.83 | 418.23 | 20 | y4 |
| | GPSVFPLAPSSK | heavy | 597.83 | 854.49 | 26 | y8 |
| | GPSVFPLAPSSK | heavy | 597.83 | 707.42 | 18 | y7 |
| | GPSVFPLAPSSK | heavy | 597.83 | 497.28 | 20 | y5 |
| | GPSVFPLAPSSK | heavy | 597.83 | 426.24 | 20 | y4 |
| | EEQYN[GlcNAc]STYR | light | 696.80 | 204.09 | 21 | GlcNAc |
| | EEQYN[GlcNAc]STYR | light | 696.80 | 843.38 | 21 | y5 |
| | EEQYN[GlcNAc]STYR | light | 696.80 | 640.19 | 21 | y5* |
| | EEQYN[GlcNAc]STYR | light | 696.80 | 526.26 | 21 | y4 |
| | EEQYN[GlcNAc]STYR | heavy | 701.80 | 204.09 | 21 | GlcNAc |
| | EEQYN[GlcNAc]STYR | heavy | 701.80 | 853.39 | 21 | y5 |
| | EEQYN[GlcNAc]STYR | heavy | 701.80 | 650.20 | 21 | y5* |
| | EEQYN[GlcNAc]STYR | heavy | 701.80 | 536.27 | 21 | y4 |
| human IgG2 | GLPAPIEK | light | 412.75 | 654.38 | 15 | y6 |
| | GLPAPIEK | light | 412.75 | 557.33 | 15 | y5 |
| | GLPAPIEK | light | 412.75 | 486.29 | 15 | y4 |
| | GLPAPIEK | light | 412.75 | 276.16 | 15 | y2 |
| | GLPAPIEK | heavy | 416.75 | 662.40 | 15 | y6 |
| | GLPAPIEK | heavy | 416.75 | 565.34 | 15 | y5 |
| | GLPAPIEK | heavy | 416.75 | 494.31 | 15 | y4 |
| | GLPAPIEK | heavy | 416.75 | 284.17 | 15 | y2 |
| | EEQFN[GlcNAc]STFR | light | 680.80 | 204.09 | 23 | GlcNAc |
| | EEQFN[GlcNAc]STFR | light | 680.80 | 974.46 | 15 | y6 |
| | EEQFN[GlcNAc]STFR | light | 680.80 | 827.39 | 15 | y5 |
| | EEQFN[GlcNAc]STFR | light | 680.80 | 510.27 | 21 | y4 |
| | EEQFN[GlcNAc]STFR | heavy | 685.81 | 204.09 | 23 | GlcNAc |
| | EEQFN[GlcNAc]STFR | heavy | 685.81 | 984.47 | 15 | y6 |
| | EEQFN[GlcNAc]STFR | heavy | 685.81 | 837.40 | 15 | y5 |
| | EEQFN[GlcNAc]STFR | heavy | 685.81 | 520.28 | 21 | y4 |
| human IgG3 | SCDTPPPCPR | light | 593.75 | 824.41 | 21 | y7 |
| | SCDTPPPCPR | light | 593.75 | 723.36 | 21 | y6 |
| | SCDTPPPCPR | light | 593.75 | 626.31 | 21 | y5 |
| | SCDTPPPCPR | light | 593.75 | 529.26 | 21 | y4 |
| | SCDTPPPCPR | heavy | 598.76 | 834.42 | 21 | y7 |
| | SCDTPPPCPR | heavy | 598.76 | 733.37 | 21 | y6 |
| | SCDTPPPCPR | heavy | 598.76 | 636.32 | 21 | y5 |
| | SCDTPPPCPR | heavy | 598.76 | 539.26 | 21 | y4 |

| | | | | | | |
|---|---|---|---|---|---|---|
| human | GLPSSIEK | light | 415.73 | 660.36 | 15 | y6 |
| IgG4 | GLPSSIEK | light | 415.73 | 563.30 | 15 | y5 |
| | GLPSSIEK | light | 415.73 | 476.27 | 15 | y4 |
| | GLPSSIEK | heavy | 419.74 | 668.37 | 15 | y6 |
| | GLPSSIEK | heavy | 419.74 | 571.32 | 15 | y5 |
| | GLPSSIEK | heavy | 419.74 | 484.29 | 15 | y4 |
| human | EEQYN[GlcNAc]STFR | light | 688.80 | 204.09 | 25 | GlcNAc |
| IgG3&4 | EEQYN[GlcNAc]STFR | light | 688.80 | 1118.51 | 14 | y7 |
| | EEQYN[GlcNAc]STFR | light | 688.80 | 915.32 | 20 | y7* |
| | EEQYN[GlcNAc]STFR | light | 688.80 | 990.45 | 16 | y6 |
| | EEQYN[GlcNAc]STFR | heavy | 693.81 | 204.09 | 25 | GlcNAc |
| | EEQYN[GlcNAc]STFR | heavy | 693.81 | 1128.52 | 14 | y7 |
| | EEQYN[GlcNAc]STFR | heavy | 693.81 | 925.33 | 20 | y7* |
| | EEQYN[GlcNAc]STFR | heavy | 693.81 | 1000.46 | 16 | y6 |
| mouse | VNSAAFPAPIEK | light | 622.34 | 872.49 | 20 | y8 |
| IgG1 | VNSAAFPAPIEK | light | 622.34 | 801.45 | 16 | y7 |
| | VNSAAFPAPIEK | light | 622.34 | 654.38 | 20 | y5 |
| | VNSAAFPAPIEK | light | 622.34 | 486.29 | 32 | y4 |
| | VNSAAFPAPIEK | heavy | 626.34 | 880.5 | 20 | y8 |
| | VNSAAFPAPIEK | heavy | 626.34 | 809.46 | 16 | y7 |
| | VNSAAFPAPIEK | heavy | 626.34 | 662.4 | 20 | y5 |
| | VNSAAFPAPIEK | heavy | 626.34 | 494.31 | 32 | y4 |
| | EEQIN[GlcNAc]STFR | light | 663.81 | 204.09 | 20 | GlcNAc |
| | EEQIN[GlcNAc]STFR | light | 663.81 | 827.39 | 22 | y5 |
| | EEQIN[GlcNAc]STFR | light | 663.81 | 624.20 | 24 | y5* |
| | EEQIN[GlcNAc]STFR | light | 663.81 | 510.27 | 24 | y4 |
| | EEQIN[GlcNAc]STFR | heavy | 668.82 | 204.09 | 20 | GlcNAc |
| | EEQIN[GlcNAc]STFR | heavy | 668.82 | 837.40 | 22 | y5 |
| | EEQIN[GlcNAc]STFR | heavy | 668.82 | 634.21 | 24 | y5* |
| | EEQIN[GlcNAc]STFR | heavy | 668.82 | 520.28 | 24 | y4 |
| rt | AGGSSEPVTGLADK | light | 644.82 | 800.45 | 22 | y8 |
| peptides | AGGSSEPVTGLADK | light | 644.82 | 604.33 | 22 | y6 |
| | VEATFGVDESANK | light | 683.83 | 966.45 | 23 | y9 |
| | VEATFGVDESANK | light | 683.83 | 819.38 | 23 | y8 |
| | YILAGVESNK | light | 547.30 | 817.44 | 19 | y8 |
| | YILAGVESNK | light | 547.30 | 633.32 | 19 | y6 |
| | TPVISGGPYYER | light | 669.84 | 1041.50 | 23 | y9 |
| | TPVISGGPYYER | light | 669.84 | 928.42 | 23 | y8 |
| | TPVITGAPYYER | light | 683.85 | 956.45 | 23 | y8 |
| | TPVITGAPYYER | light | 683.85 | 855.40 | 23 | y7 |
| | GDLDAASYYAPVR | light | 699.34 | 926.47 | 24 | y8 |
| | GDLDAASYYAPVR | light | 699.34 | 855.44 | 24 | y7 |
| | TGFIIDPGGVIR | light | 622.85 | 713.39 | 22 | y7 |
| | TGFIIDPGGVIR | light | 622.85 | 598.37 | 22 | y6 |
| | GTFIIDPAAIVR | light | 636.87 | 854.51 | 22 | y8 |
| | GTFIIDPAAIVR | light | 636.87 | 626.40 | 22 | y6 |
| | ADVTPADFSEWSK | light | 726.84 | 1066.48 | 25 | y9 |
| | ADVTPADFSEWSK | light | 726.84 | 387.19 | 25 | b4 |

TABLE S3: SRM DETECTION LIMITS AND CONVERSION FACTORS

Lower limits of quantification (LLoQ) and conversion factors are given for all the peptides analyzed by SRM in this study. The conversion factor denotes the ratio between the measured amounts and the amounts of each peptide spiked in.

| Protein | Peptide sequence | LLoQ (fmol) | Conversion factor |
|---|---|---|---|
| human IgG1 | GPSVFPLAPSSK | 1.62 | 1.74 |
| | EEQYN[GlcNAc]STYR | 6.41 | 2.15 |
| human IgG2 | GLPAPIEK | 1.45 | 1.76 |
| | EEQFN[GlcNAc]STFR | 1.98 | 1.23 |
| human IgG3 | SCDTPPPCPR | 0.16 | 1.64 |
| human IgG4 | GLPSSIEK | 2.38 | 1.72 |
| human IgG3&4 | EEQYN[GlcNAc]STFR | 1.19 | 1.74 |
| mouse IgG1 | VNSAAFPAPIEK | 1.11 | 1.29 |
| | EEQIN[GlcNAc]STFR | 0.96 | 3.47 |

Table S4: Tonsillitis Patients

See excel sheet

Table S5: Sepsis Patients

See excel sheet

Table S6: Statistical analysis of IgG glycan hydorlysis in tonsillitis patients

| | GAS-negative | GAS positive |
|---|---|---|
| Number of values (n) | 28 | 26 |
| Median | 0 | 10.3 |
| Mean | 3.896 | 14.98 |
| 95% CI | 0.418-7.374 | 7.389-22.56 |

| Mann-Whitney test | |
|---|---|
| p value | 0.0057 |
| Tails | Two-tailed |
| Mann-Whitney U | 223.5 |

Table S7: Analysis of correlation between IgG glycan hydorlysis and disease parameters

| | General malaise | Throat pain | Centor score |
|---|---|---|---|
| Number of values (n) | 26 | 26 | 26 |
| Spearman ρ | 0.5036 | 0.4583 | 0.4406 |
| 95% CI | 0.1326-0.7508 | 0.0743-0.724 | 0.05222-0.7133 |
| p value | 0.0087 | 0.0185 | 0.0243 |
| Tails | Two-tailed | Two-tailed | Two-tailed |

TABLE S8: STATISTICAL ANALYSIS OF IGG LEVELS IN SEPSIS PATIENT PLASMA

| | healthy | sepsis | Severe sepsis | Septic shock | GAS sepsis | GAS severe sepsis | GAS septic shock |
|---|---|---|---|---|---|---|---|
| Number of values | 12 | 4 | 6 | 4 | 3 | 9 | 6 |
| Median | 12.57 | 11.44 | 9.811 | 7.849 | 7.698 | 9.451 | 12.07 |
| Mean | 12.42 | 11.41 | 9.133 | 8.054 | 7.088 | 8.837 | 14.11 |
| 95% CI | 10.07-14.78 | 5.782-17.04 | 7.401-10.86 | 3.929-12.18 | 4.15710.02 | 7.47-10.2 | 3.661-24.55 |

| Kruskal-Wallis test | |
|---|---|
| P value | 0.0622 |
| Kruskal-Wallis statistic | 11.99 |
| Tails | Two-tailed |
| **Dunn's multiple comparisons test** | **Adjusted P Value** |
| healthy vs. sepsis | >0.9999 |
| healthy vs. severe sepsis | >0.9999 |
| healthy vs. septic shock | 0.7689 |
| healthy vs. GAS sepsis | 0.2118 |
| healthy vs. GAS severe sepsis | 0.5056 |
| healthy vs. GAS septic shock | >0.9999 |
| sepsis vs. severe sepsis | >0.9999 |
| sepsis vs. septic shock | >0.9999 |
| sepsis vs. GAS sepsis | >0.9999 |
| sepsis vs. GAS severe sepsis | >0.9999 |
| sepsis vs. GAS septic shock | >0.9999 |
| severe sepsis vs. septic shock | >0.9999 |
| severe sepsis vs. GAS sepsis | >0.9999 |
| severe sepsis vs. GAS severe sepsis | >0.9999 |
| severe sepsis vs. GAS septic shock | >0.9999 |
| septic shock vs. GAS sepsis | >0.9999 |
| septic shock vs. GAS severe sepsis | >0.9999 |
| septic shock vs. GAS septic shock | >0.9999 |
| GAS sepsis vs. GAS severe sepsis | >0.9999 |
| GAS sepsis vs. GAS septic shock | 0.9977 |
| GAS severe sepsis vs. GAS septic shock | >0.9999 |

TABLE S9: STATISTICAL ANALYSIS OF IGG GLCYAN HYDROLYSIS IN SEPSIS PATIENT PLASMA

| | healthy | sepsis | Severe sepsis | Septic shock | GAS sepsis | GAS severe sepsis | GAS septic shock |
|---|---|---|---|---|---|---|---|
| Number of values | 12 | 4 | 6 | 4 | 3 | 9 | 6 |
| Median | 0 | 0 | 0 | 0 | 0 | 0 | 0.091 |
| Mean | 0 | 0 | 0 | 0 | 0 | 0 | 0.25 |
| 95% CI | 0 | 0 | 0 | 0 | 0 | 0 | -0.15-0.66 |

| Kruskal-Wallis test | |
|---|---|
| P value | <0.0001 |
| Kruskal-Wallis statistic | 35 |
| Tails | Two-tailed |

| Dunn's multiple comparisons test | Adjusted P Value |
|---|---|
| healthy vs. sepsis | >0.9999 |
| healthy vs. severe sepsis | >0.9999 |
| healthy vs. septic shock | >0.9999 |
| healthy vs. GAS sepsis | >0.9999 |
| healthy vs. GAS severe sepsis | >0.9999 |
| healthy vs. GAS septic shock | <0.0001 |
| sepsis vs. severe sepsis | >0.9999 |
| sepsis vs. septic shock | >0.9999 |
| sepsis vs. GAS sepsis | >0.9999 |
| sepsis vs. GAS severe sepsis | >0.9999 |
| sepsis vs. GAS septic shock | 0.0013 |
| severe sepsis vs. septic shock | >0.9999 |
| severe sepsis vs. GAS sepsis | >0.9999 |
| severe sepsis vs. GAS severe sepsis | >0.9999 |
| severe sepsis vs. GAS septic shock | 0.0002 |
| septic shock vs. GAS sepsis | >0.9999 |
| septic shock vs. GAS severe sepsis | >0.9999 |
| septic shock vs. GAS septic shock | 0.0013 |
| GAS sepsis vs. GAS severe sepsis | >0.9999 |
| GAS sepsis vs. GAS septic shock | 0.0052 |
| GAS severe sepsis vs. GAS septic shock | <0.0001 |

TABLE S10: STATISTICAL ANALYSIS OF PHAGOCYTIC KILLING – MACROPHAGES HIGH ANTI-M1

| | 5448 | 5448+rEndoS | 5448ndoS | 5448ndoS +rEndoS |
|---|---|---|---|---|
| Number of values | 9 | 9 | 9 | 9 |
| Median | 81.43 | 84.3 | 61.14 | 87.5 |
| Mean | 89.9 | 84.35 | 57.59 | 91.66 |
| 95% CI | 72.53-107.3 | 74.81-93.89 | 48.64-66.53 | 71.19-112.1 |

| ANOVA summary | |
|---|---|
| F | 5.98 |
| P value | 0.0023 |
| Tails | Two-tailed |
| Tukey's multiple comparisons test | Adjusted P Value |
| 5448 vs. 5448+rEndoS | 0.9293 |
| 5448 vs. 5448ndoS | 0.0067 |
| 5448 vs. 5448ndoS + rEndoS | 0.9974 |
| 5448+rEndoS vs. 5448ndoS | 0.0306 |
| 5448+rEndoS vs. 5448ndoS | 0.8547 |
| 5448+rEndoS vs. 5448ndoS + rEndoS | 0.004 |
| 5448ndoS vs. 5448ndoS + rEndoS | 0.9293 |

TABLE S11: STATISTICAL ANALYSIS OF PHAGOCYTIC KILLING – MACROPHAGES LOW ANTI-M1

| | 5448 | 5448+rEndoS | 5448ndoS | 5448ndoS +rEndoS |
|---|---|---|---|---|
| Number of values | 9 | 9 | 9 | 9 |
| Median | 88.68 | 86.25 | 97.63 | 96.67 |
| Mean | 87.48 | 87.32 | 93.52 | 97.62 |
| 95% CI | 77.98-96.98 | 79.57-95.08 | 80.73-106.3 | 85.39-109.9 |

| ANOVA summary | |
|---|---|
| F | 1.15 |
| P value | 0.3441 |
| Tails | Two-tailed |
| Tukey's multiple comparisons test | Adjusted P Value |
| 5448 vs. 5448+rEndoS | >0.9999 |
| 5448 vs. 5448ndoS | 0.797 |
| 5448 vs. 5448ndoS + rEndoS | 0.4284 |
| 5448+rEndoS vs. 5448ndoS | 0.7845 |
| 5448+rEndoS vs. 5448ndoS | 0.4151 |
| 5448+rEndoS vs. 5448ndoS + rEndoS | 0.9245 |
| 5448ndoS vs. 5448ndoS + rEndoS | >0.9999 |

TABLE S12: STATISTICAL ANALYSIS OF PHAGOCYTIC KILLING – POLYMORPHONUCLEAR CELLS HIGH ANTI-M1

| | 5448 | 5448+rEndoS | 5448ndoS | 5448ndoS +rEndoS |
|---|---|---|---|---|
| Number of values | 9 | 9 | 9 | 9 |
| Median | 112.5 | 108.5 | 74.7 | 100.5 |
| Mean | 105.9 | 105.6 | 68.83 | 106.8 |
| 95% CI | 93.78-118 | 86.29-124.9 | 53.24-84.43 | 91.28-122.3 |

| ANOVA summary | |
|---|---|
| F | 7.357 |
| P value | 0.0007 |
| Tails | Two-tailed |
| Tukey's multiple comparisons test | Adjusted P Value |
| 5448 vs. 5448+rEndoS | >0.9999 |
| 5448 vs. 5448ndoS | 0.0032 |
| 5448 vs. 5448ndoS + rEndoS | 0.9997 |
| 5448+rEndoS vs. 5448ndoS | 0.0034 |
| 5448+rEndoS vs. 5448ndoS | 0.9993 |
| 5448+rEndoS vs. 5448ndoS + rEndoS | 0.0024 |
| 5448ndoS vs. 5448ndoS + rEndoS | >0.9999 |

Table S13: Statistical analysis of phagocytic killing – Polymorphonuclear cells low anti-M1

| | 5448 | 5448+rEndoS | 5448ndoS | 5448ndoS +rEndoS |
|---|---|---|---|---|
| Number of values | 9 | 9 | 9 | 9 |
| Median | 88.72 | 103.8 | 98.08 | 104.3 |
| Mean | 95.83 | 103.6 | 95.32 | 101.8 |
| 95% CI | 81.95-109.7 | 92.4-114.7 | 85.44-105.2 | 90.56-113.0 |

| ANOVA summary | |
|---|---|
| F | 0.6821 |
| P value | 0.5695 |
| Tails | Two-tailed |
| **Tukey's multiple comparisons test** | **Adjusted P Value** |
| 5448 vs. 5448+rEndoS | 0.7014 |
| 5448 vs. 5448ndoS | 0.9999 |
| 5448 vs. 5448ndoS + rEndoS | 0.8388 |
| 5448+rEndoS vs. 5448ndoS | 0.6579 |
| 5448+rEndoS vs. 5448ndoS | 0.9943 |
| 5448+rEndoS vs. 5448ndoS + rEndoS | 0.8026 |
| 5448ndoS vs. 5448ndoS + rEndoS | 0.7014 |

Figure S1: IgG SRM assay calibration – human IgG subclasses

Measured amounts of each peptide are plotted against the theoretical amounts. The line and shading mark measurements below the limit of quantification.

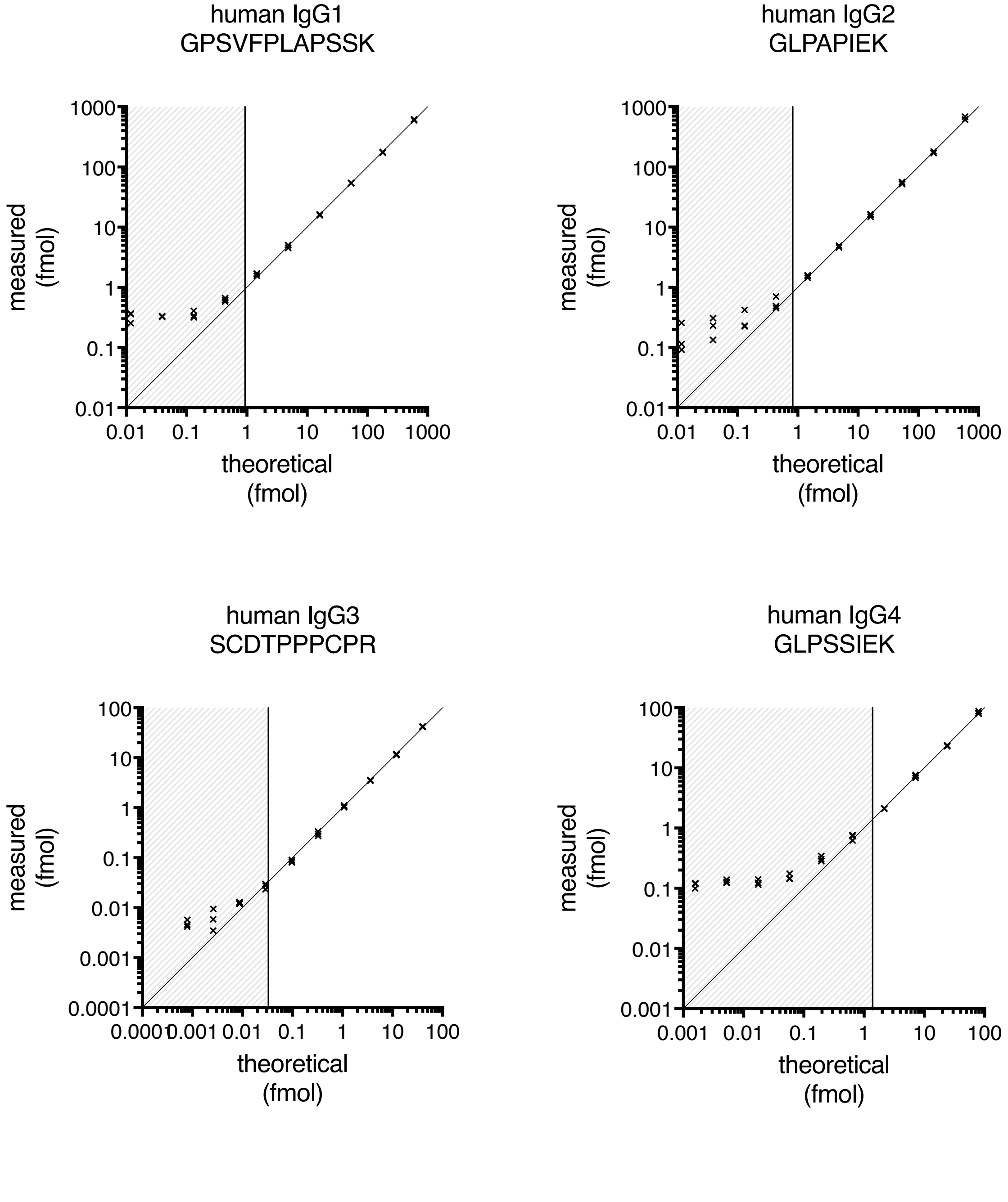

FIGURE S2: IGG SRM ASSAY CALIBRATION – HUMAN IGG GLYCOPEPTIDES

Measured amounts of each peptide are plotted against the theoretical amounts. The line and shading mark measurements below the limit of quantification.

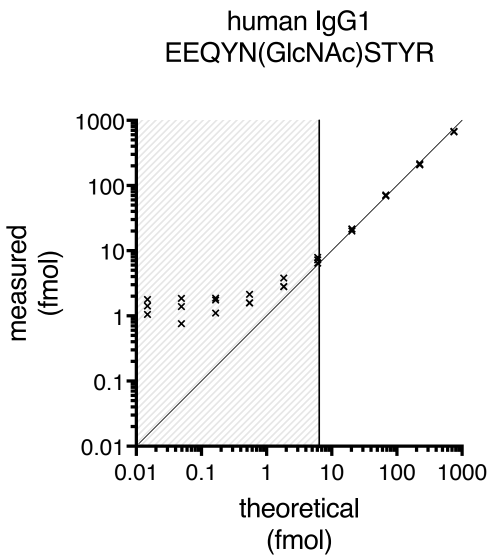


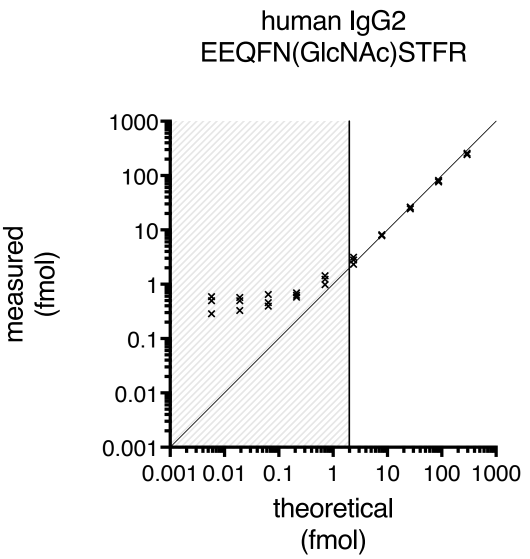


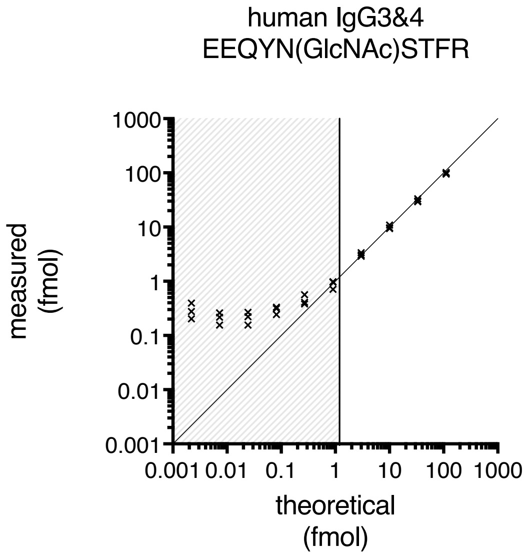

FIGURE S3: ANTI-M1 IGG RESPONSE IN DONOR SERA FOR PHAGOCYTOSIS ASSAYS

IgG response to MA was determined by ELISA and signal in respect to serum dilution is shown. Donors 1-3 were classified as high anti-M1 response, Donor 4 as low anti-M1 response. Normal mouse serum (NMS) acted as negative control.

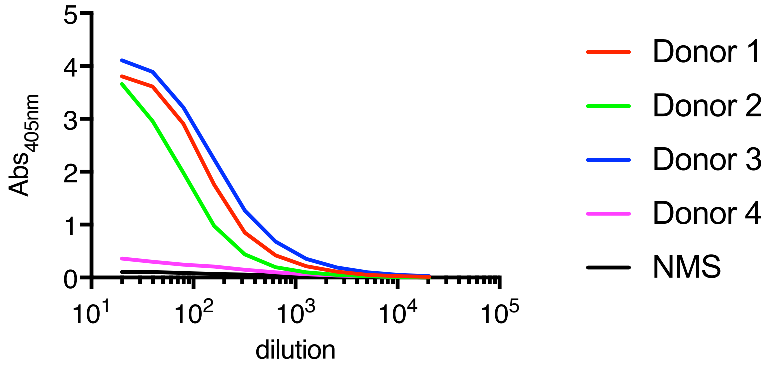

FIGURE S4: IGG SRM ASSAY CALIBRATION – MURINE IGG1

Measured amounts of each peptide are plotted against the theoretical amounts. The line and shading mark measurements below the limit of quantification.

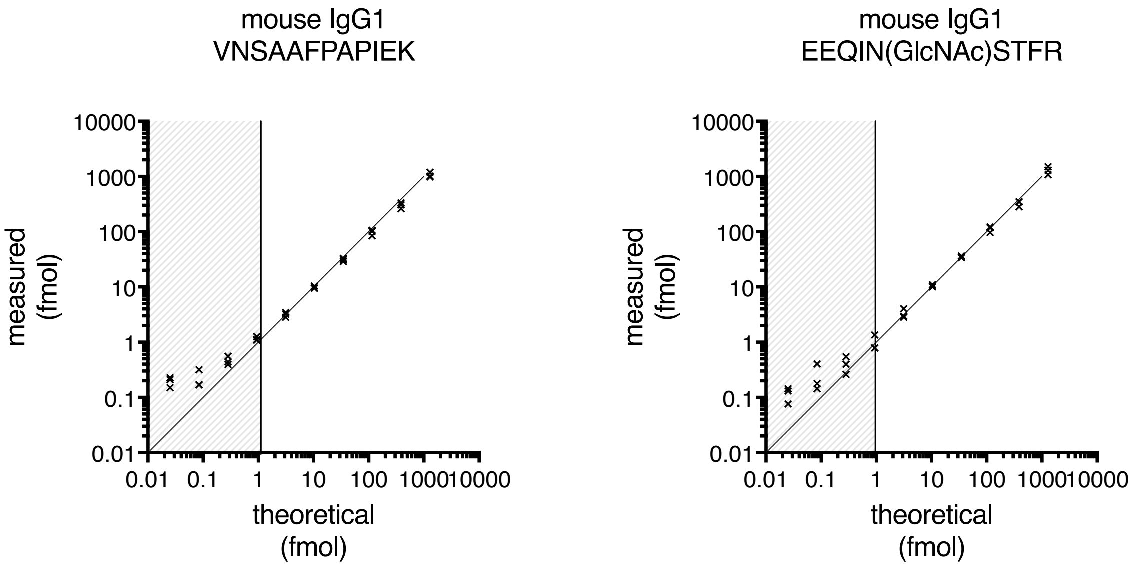

FIGURE S5: CORRELATION BETWEEN SKIN BACTERIAL LOAD AND ABSOLUTE AMOUNT OF $IgG_{GH}$

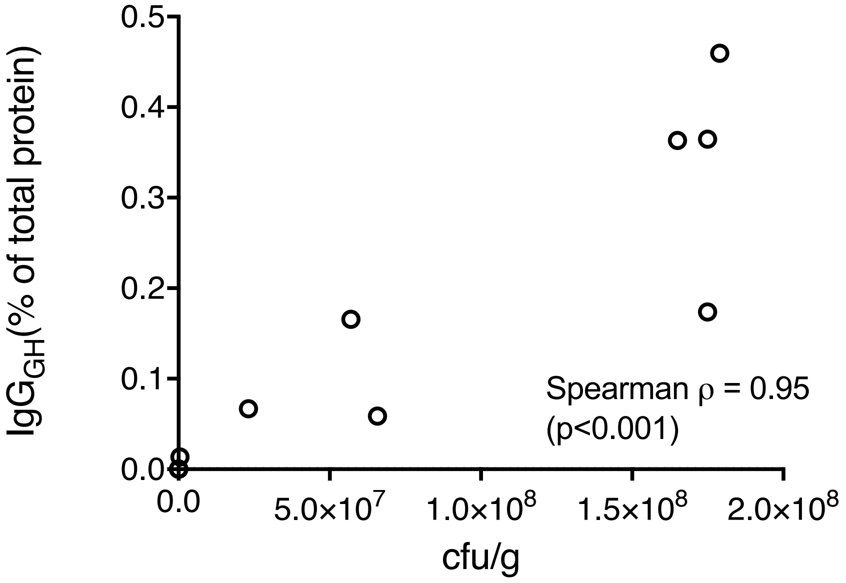

FIGURE S6: IGG DEGRADATION BY GAS AP1 GROWN IN SALIVA

GAS strains AP1, 5448, and their respective *ndoS* mutants were grown in human saliva supplemented with 5% serum and the culture supernatants were analyzed by SDS-PAGE under non-reducing conditions. Note the lack of fully intact IgGs in the AP1 culture supernatants.

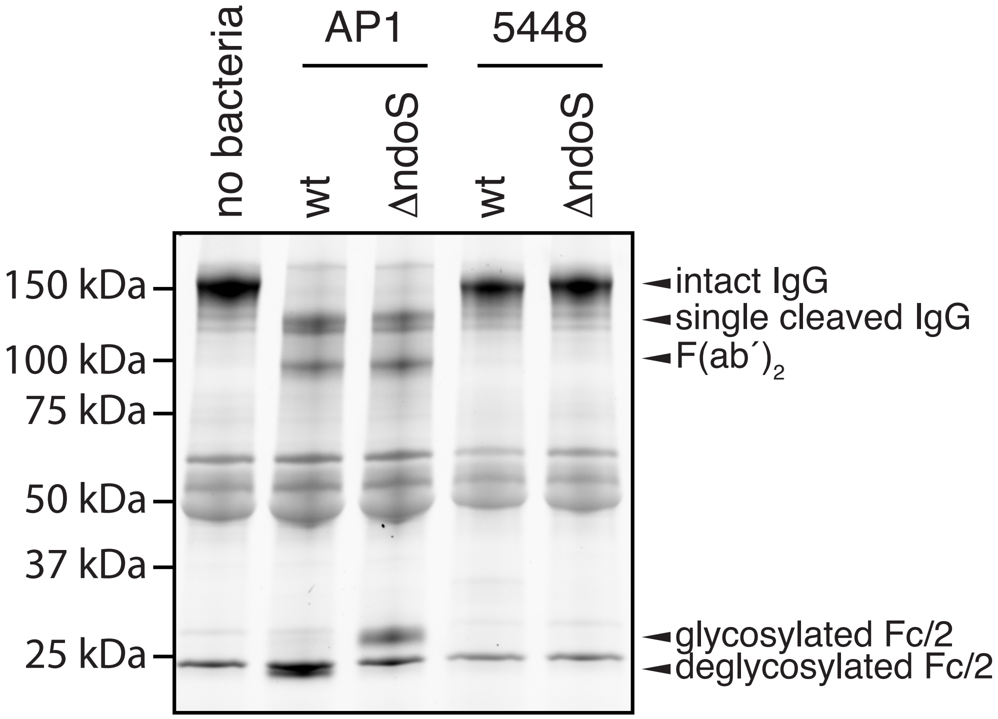

### Figure S7: subcutaneous infection of naïve mice – Disease parameters

Bacterial loads in skin and spleen as well as loss of body weight were determined 48h post infection. Data was analyzed using a Kruskal-Wallis test followed by Dunn´s multiple comparison test (*: $p<0.05$)

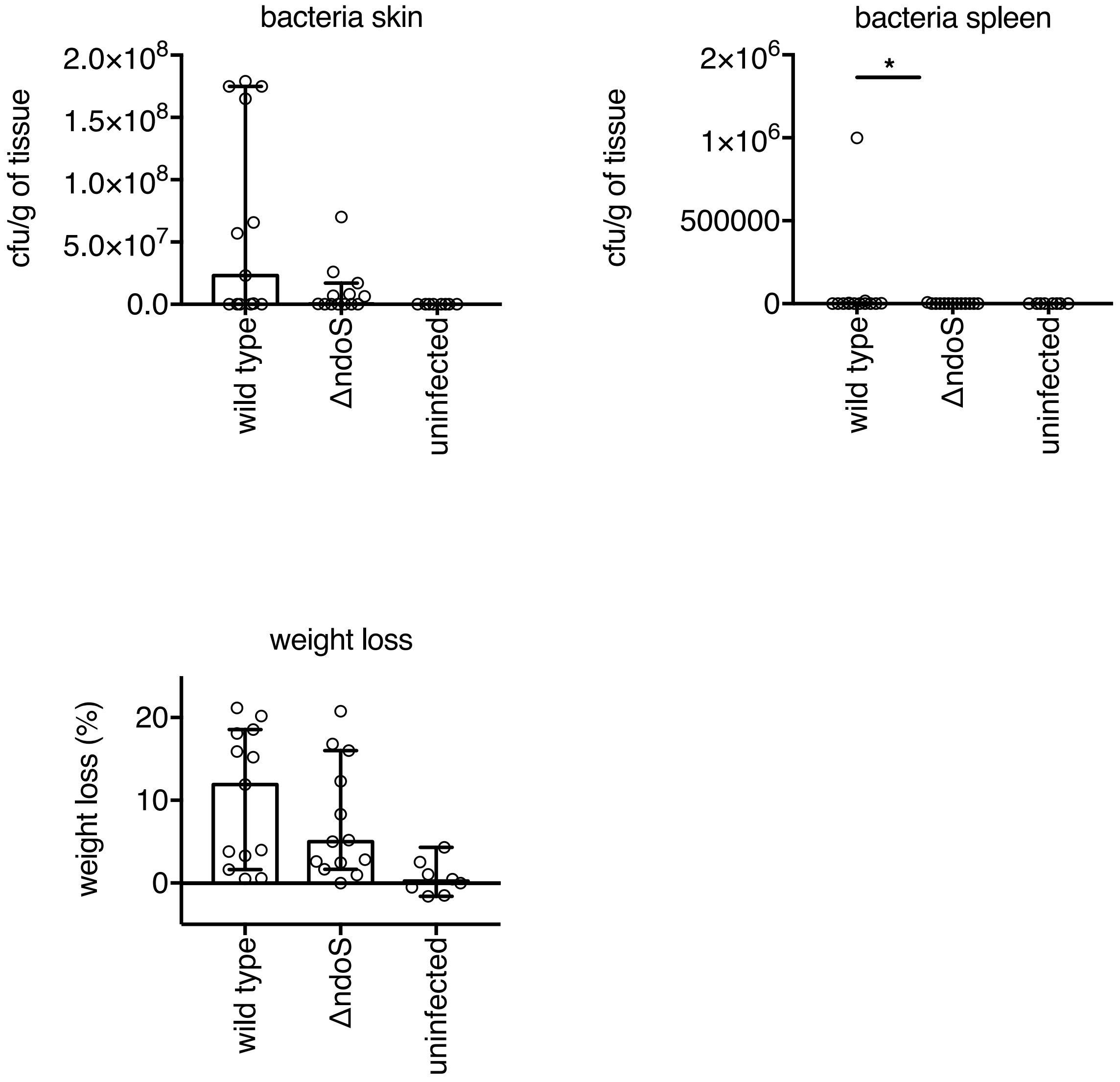

FIGURE S8: SUBCUTANEOUS INFECTION OF NAÏVE MICE – IGG GLYCAN HYDROLYSIS

Glycan hydrolysis of IgG1 was determined using SRM mass spectrometry in skin homogenates and plasma samples taken at 48h post infection.

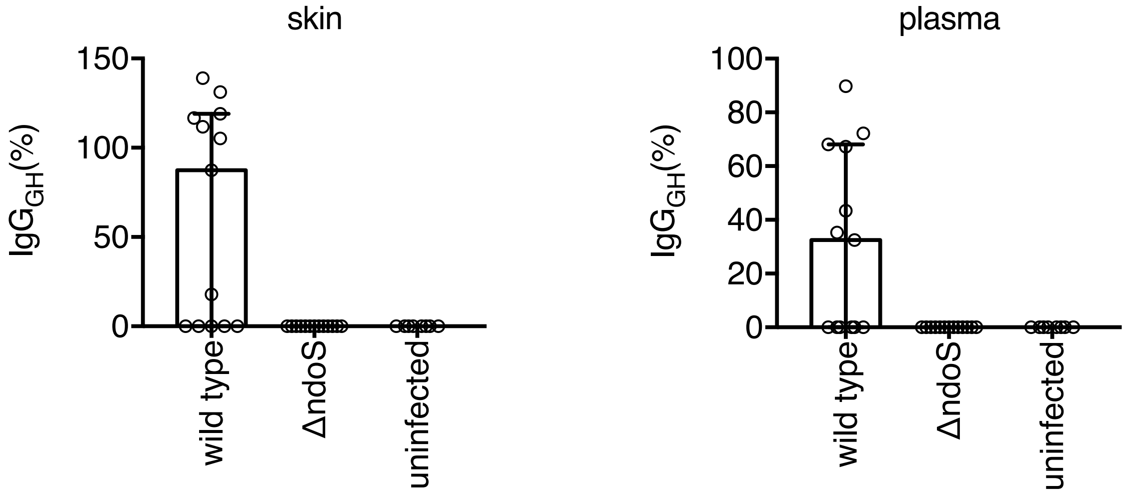

FIGURE S9: IMMUNIZATION WITH M1 PROTECTS AGAINST GAS INFECTION

Mice were immunized by two injections with M1 protein (red lines) or mock-immunized with adjuvant only (black line). 5-day survival was monitored after infection with GAS AP1 at a low dose ($2.5 \times 10^5$ cfu, unbroken lines) or a high dose ($2 \times 10^7$ cfu, dashed line).

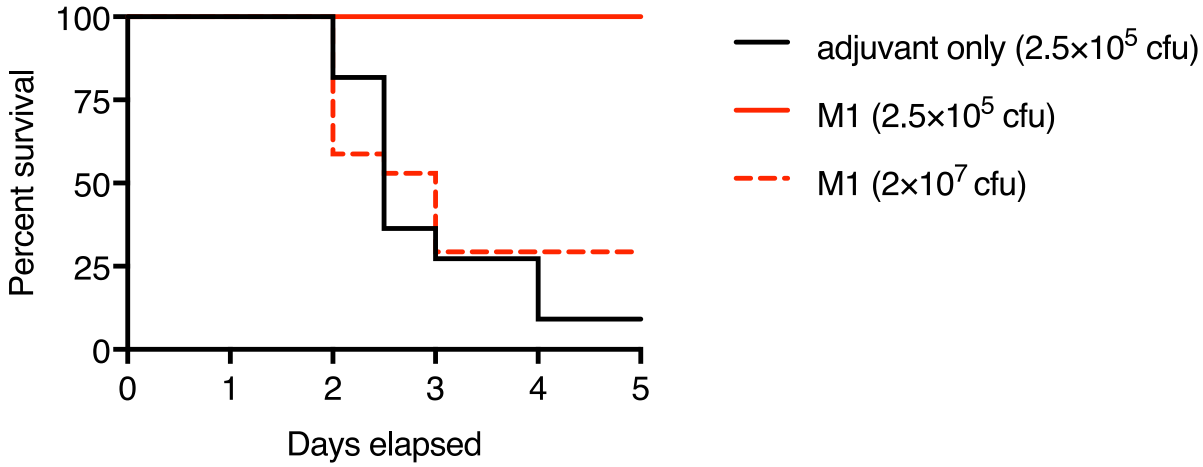

FIGURE S9: CHANGES IN BODY WEIGHT DURING GAS INFECTION IN MICE

Changes in body weight of mice infected with AP1 wild type (black) or *ndo*S mutant bacteria (red line). Each line depicts an individual animal.

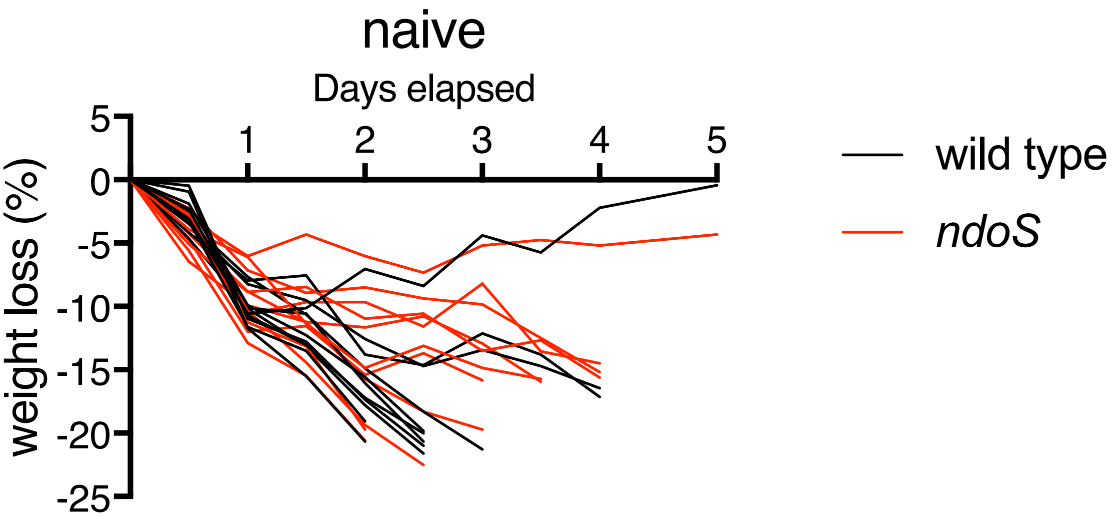


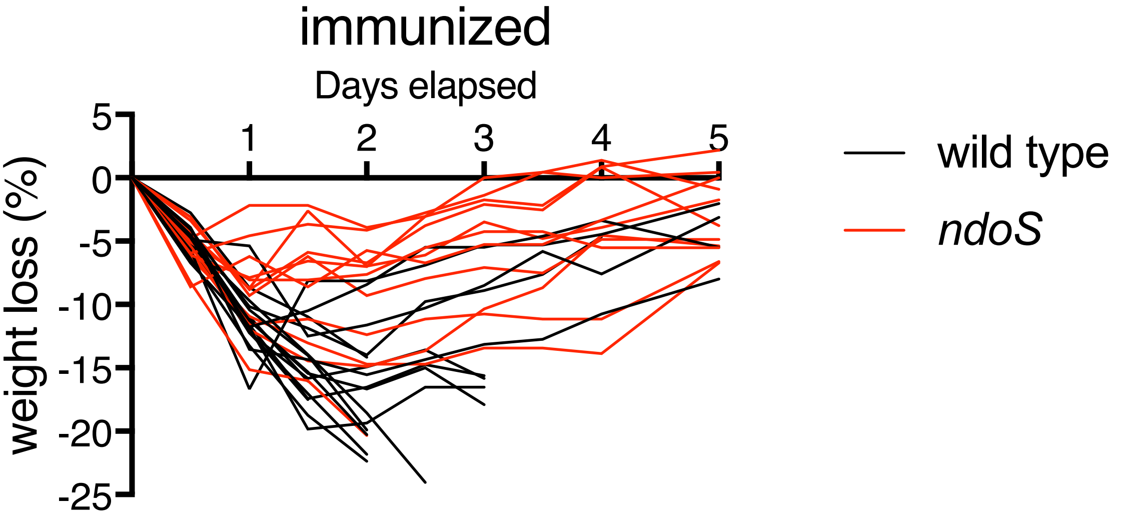
